# Simulating the spectrum, not the syndrome: Large scale individualized modeling of oral reading in stroke aphasia

**DOI:** 10.64898/2026.05.11.724319

**Authors:** Ryan Staples, Andrew T. DeMarco, Alycia B. Laks, Peter E. Turkeltaub

**Author notes:** Emails: Ryan Staples, Andrew T. DeMarco, Alycia B. Laks, Peter E. Turkeltaub. Correspondence to: Ryan Staples, 4000 Reservoir Rd NW, Building D, Suite 165, Washington, DC 20057.

## Abstract

Computational models are a linchpin in our understanding of the neurocognitive basis of reading. These models can simulate idealized profiles of alexia syndromes, but in reality, individuals with alexia present with a wide range of mixed deficits rather than idealized syndromes. To provide a complete cognitive theory of reading, computational models must be able to account for this individual variation. However, this has never been demonstrated. We test oral reading and non-reading phonological and semantic processing in 83 left-hemisphere stroke survivors. We show that individual alexia profiles can be simulated by applying graded phonology and semantic lesions to an artificial neural network model of reading, creating “matched models” that represent individual stroke survivors. The severity of damage to the semantic and phonological layers of the matched models was highly correlated with directly-measured semantic and phonological processing deficits. However, we also identify systematic ways in which the models fail to simulate the reading performance of their matched stroke survivors. Our results support theories of alexia that rely on process-based deficits, demonstrate the feasibility of large-scale individualized modelling of alexia, and suggest ways to further improve the correspondence of models and human reading behavior.

## Introduction

Computational models have provided a means of instantiating quantitative hypotheses about cognitive architecture and processing. One domain in which computational modelling has a long history is language processing, and specifically in reading. Computational models have been used to simulate many phenomena in typical reading (Coltheart, Rastle, Perry, Langdon, & Ziegler, 2001; Perry, Ziegler, & Zorzi, 2007; Seidenberg & McClelland, 1989) and the profiles of prototypical cases of acquired reading disorders, i.e., alexias (Plaut, McClelland, Seidenberg, & Patterson, 1996; Welbourne, Woollams, Crisp, & Lambon Ralph, 2011; Woollams, Lambon Ralph, Plaut, & Patterson, 2007). In doing so, they have revealed a great deal about the cognitive architecture of reading. Still, many individual cases of alexia often do not exhibit the prototypical patterns that have been simulated by these models (Brookshire, Wilson, Nadeau, Rothi, & Kendall, 2014; Ripamonti et al., 2014). Prominent computational models of picture naming and repetition have been shown to predict the wide variation in individual cases of aphasia, simultaneously validating the models and pointing toward ways to improve them (Dell, Martin, & Schwartz, 2007; Foygel & Dell, 2000; Schwartz, Dell, Martin, Gahl, & Sobel, 2006).

Here, we aim to do the same for a prominent model of reading, leveraging a large sample of left-hemisphere stroke survivors with varying degrees of reading and cognitive impairments. We utilize artificial neural networks, which allow for simulations of graded impairment to the underlying cognitive processes that support reading, to move away from the traditional syndrome-based approach. In doing so, we provide a strict test of the capacity of the models to accommodate a full spectrum of individual variability in reading impairment and identify where the models fail to capture reading and thus may be improved.

### Syndromic classifications of reading disorders

Alexia, an acquired reading disorder, affects two in every three stroke survivors with aphasia (Brookshire, Wilson, et al., 2014). Reflecting the importance of reading in contemporary societies, this loss of literacy in post-stroke alexia is experienced as a serious negative impact on quality of life, and persons with alexia express a strong desire to improve their reading ability (Madden & Bush, 2024). However, while reading often recovers to some extent, recovery is highly variable and usually incomplete (Bahrami Balani & Bickerton, 2023; Friedman, 1996; Newcombe, Marshall, Carrivick, & Hiorns, 1974).

Alexias have traditionally been classified into syndromes, reflecting constellations of impairments, error types, and compensatory strategies. Two alexia syndromes, phonological and surface alexia, have been prominent in the development of theories of reading. Phonological alexia describes a condition in which the reading of familiar words is relatively preserved, but the reading of unfamiliar or pseudowords (graphotactically correct, pronounceable letter strings) is impaired (Beauvois & Derouesne, 1979; Derouesne & Beauvois, 1979). Surface alexia, on the other hand, is characterized by the relatively preserved reading of consistent words (words that follow the most common spelling-to-sound correspondences) and pseudowords but impaired reading of inconsistent words, particularly low frequency inconsistent words (Coltheart, Masterson, Byng, Prior, & Riddoch, 1983). Many of the errors made on low frequency words are “regularization” or “legitimate alternate reading of component” (LARC) errors, where the word is pronounced as if it were consistent (e.g. “yacht”➔ /jælllt/) (Patterson, Marshall, & Colthart, 1985; Woollams et al., 2007).

### Cognitive models of oral reading

The dissociation between phonological and surface alexia suggests that there are at least two cognitive processes that interact to support reading: one mechanism that supports whole-word “lexical” reading, allowing reading of known words, and a second that decodes sound from print, allowing “sublexical” reading of pseudowords. This two-route structure is present in the two most prominent cognitive models of reading. The Dual-Route Cascaded (DRC) model implements the lexical pathway using localist representations for each word that the model “knows”. Visual features activate positional letter units, which then excite all word units in an orthographic lexicon that have that letter in the appropriate position and inhibit any that do not. This activation is transferred to a phonological lexicon unit, which then excites phoneme units. The sublexical pathway in the DRC model is implemented using either a set of symbolic rules (Coltheart et al., 2001) or, in the connectionist dual-process plus (CDP+) model, a two-layer artificial neural network (ANN) to decode sound from spelling (Perry et al., 2007). DRC simulations of phonological alexia are performed by impairing the function of the sublexical pathway, leading to a reliance on whole-word processing via the lexicon. Simulations of surface alexia impair the function of the lexicons, forcing a reliance on the sublexical pathway (Woollams, Lambon Ralph, Plaut, & Patterson, 2010). However, the DRC does not have a learning algorithm, making it incapable of simulating learning or recovery from alexia.

An alternative to the DRC proposes that the mechanisms that support reading are not specific to reading. The primary systems hypothesis (PSH) argues that reading relies on generalized systems that support vision, phonology, and semantic processing (Patterson & Lambon Ralph, 1999; Woollams, 2014). The PSH was motivated by the observation that alexia is broadly accompanied by generalized disorders of visual, phonological, or semantic processing (Crisp & Lambon Ralph, 2006; Roberts et al., 2013; Woollams et al., 2007). PSH models of alexia are implemented using ANN “triangle” models, which learn mappings between prespecified representations of orthography, semantics, and phonology (Plaut et al., 1996; Seidenberg & McClelland, 1989). Simulations of phonological alexia are accomplished by removing connections into or out of the phonological layer of the model, whereas simulations of surface alexia are accomplished by removing connections into or out of the semantic layer (Welbourne & Lambon Ralph, 2007; Welbourne et al., 2011; Woollams et al., 2007). Because ANN models use a learning algorithm, they are appropriate for simulating recovery from alexia. In fact, this capacity to reorganize and optimize use of the remaining computational resources is key to simulating the behavioral effects observed in alexia (Welbourne & Lambon Ralph, 2005; Welbourne et al., 2011). Plasticity-related recovery after stroke may reflect a similar reorganization of cortical resources.

While models have been successful at simulating patterns observed in idealized alexia syndromes, in reality there is great variability in the scope and severity of reading deficits in individual stroke survivors. Many stroke survivors do not exhibit patterns that can be easily classified into a classic alexia syndrome. For example, 15 of the 59 stroke survivors studied by Ripamonti et al. (2014) were classified as “undifferentiated”. In addition, a wide range of reading ability can exist within a given classification (Brookshire, Wilson, et al., 2014; Dickens et al., 2021; Ripamonti et al., 2014). Adequate models of alexia should be capable of simulating the full range of reading deficit patterns observed in individuals, both as a test of the underlying theory that they implement and so that valid inferences about rehabilitation can be drawn from the models. Here, we aim to test if a triangle model of reading can do so.

Analogous attempts to simulate a spectrum of language impairments have benefited the scientific understanding of aphasia. A series of studies combined case-series and computational approaches to evaluate and improve a model of speech errors in stroke survivors (Dell et al., 2007; Dell, Schwartz, Martin, Saffran, & Gagnon, 1997; Foygel & Dell, 2000; Schwartz et al., 2006). These studies simulated the naming performance of large samples of stroke survivors to identify where the models did and did not predict behavior. In doing so, they were able to identify that a version of their model where parameters affecting semantic and phonological processing outperformed a version that altered global parameters (Dell et al., 2007; Schwartz et al., 2006). Their results provided a critical validation of process-based theories of speech production.

### The present work

No work to our knowledge has extended Dell and Schwartz’s pioneering work modeling individual speech behavior in individuals with aphasia to assess reading behavior in alexia. Here, we assess oral reading, as well as the degree of impairment of phonological and semantic processing, in a large sample of stroke survivors with alexia. We then build individualized in silico simulations of each stroke survivor’s reading system, by damaging the phonological and semantic portions of ANN models of reading to best match each individual’s word and pseudoword reading. We then test if these individualized models recapitulate person-by-person reading profiles as well as their non-reading phonological and semantic deficits. Finally, we identified where and why the matched models poorly predict reading accuracy.

## Methods

### Participants

Participants included 83 left-hemisphere stroke survivors and 62 age- and education-matched neurotypical controls (Table 1), recruited as part of an ongoing study on individual differences in aphasia after left-hemisphere stroke (https://clinicaltrials.gov NCT04991519). Participants were native English speakers with adequate vision and hearing, corrected with lenses or hearing aids, to complete auditory and visual tasks. Controls were excluded if they scored below the cutoff adjusted for their age, ethnicity, and race (Milani, Marsiske, Cottler, Chen, & Striley, 2018) on the Montreal Cognitive Assessment (Nasreddine et al., 2005). Exclusion criteria for both the stroke and control cohorts included: history of significant neurological disease, history of head injury causing loss of consciousness, history of psychiatric disorder requiring hospitalization, ongoing use of psychiatric medications other than common antidepressants, and history of learning disorder requiring educational intervention (e.g., developmental dyslexia). All participants in the stroke cohort were at least six months post-stroke at the time of enrollment. Finally, because the model has no capacity to simulate motor speech disorders, we removed 12 stroke survivors who had severe apraxia of speech per the Apraxia of Speech Ratings Scale 3.0 (see below) (Strand, Duffy, Clark, & Josephs, 2014; Wambaugh, Bailey, Mauszycki, & Bunker, 2019). The participants consisted of all the controls and stroke survivors from within the recruitment period who fulfilled these criteria. Four controls were excluded in order to achieve demographic matching between groups. Written informed consent was obtained from all participants as required by the Declaration of Helsinki. The study protocol was approved by the Georgetown University Institutional Review Board.

**Table 1.** Sample demographics.

|  | Age,<br>years | Education,<br>by degree |  | Sex | Lesion<br>Volume<br>(cm<sup>3</sup>) | Stroke<br>Chronicity<br>(months) | WAB<br>AQ | ASRS3<br>Total of<br>Ratings |
| --- | --- | --- | --- | --- | --- | --- | --- | --- |
| Control<br>(*n* = 62) | 69.8<br>(11.6) | 17.2<br>(2.3) |  | 31F<br>31M | - | - | - | - |
| Stroke<br>(*n* = 83) | 60.5<br>(12.0) | 16.3<br>(3.0) |  | 39F<br>44M | 90.2<br>(95.8) | 49.4<br>(58.9) | 80.9<br>(18.3) | 7.7<br>(7.3) |
| U | 2507 | 2951 | X<sup>2</sup><br>(d.f.) | 0.1<br>(1) | - | - | - | - |
| *p* | 0.79 | 0.12 |  | 0.72 |  |  |  |  |

WAB AQ = Western Aphasia Battery Aphasia Quotient, ASRS3 = Apraxia of Speech Rating Scale 3.0

### Behavioral Tests

#### Oral word reading

Participants read aloud 200 monosyllabic English words, varied in word frequency (low versus high frequency of occurrence in English, such as *sheen* versus *lunch*), spelling-to-sound consistency (consistent versus inconsistent spelling-to-sound correspondences, such as *steam* vs *steak*), and imageability (Strain & Herdman, 1999; Strain, Patterson, & Seidenberg, 1995) (low versus high mental imagery, such as *grace* versus *horse*). The word list consisted of 100 words within each level of each factor. Letter length ranged from three to six letters. Low and high imageability words were matched on letter length, frequency, consistency, and articulatory complexity (Stoel-Gammon, 2010). Matching between word types was assessed through Welch’s two-sample t-tests. Briefly, each word was displayed in a pseudorandomized order in the center of a touchscreen. Participants were instructed to read aloud the word as quickly and accurately as possible within a ten-second limit. If the participant did not respond within ten seconds, the trial timed out, and the participant was prompted to proceed to the next item. Accuracy was scored for the first complete attempt, defined as the first response consisting of a consonant and vowel (CV or VC), excluding a schwa /_ə_/ (Roach, Schwartz, Martin, Grewal, & Brecher, 1996). An isolated vowel was accepted as the first complete attempt if the target itself was a CV or VC. Words were displayed in blocks of 25 items, with self-paced breaks in between blocks. Further details of the stimuli can be found in Staples et al., (2025a).

#### Oral pseudoword reading

Participants also read aloud a list of 60 pseudowords. Three types of pseudowords were presented (20 stimuli each). These 20-item pseudoword subsets were differentiated based on the number of orthography-to-phonology body mappings that exist in English: zero mappings (0M), one mapping (1M), and multiple mappings (MM) (See Dyslin et al., 2025 for details). 0M pseudowords (e.g., “dofe”) are pseudowords whose orthographic bodies do not exist within the English lexicon. 1M pseudowords (e.g., “bink”) have only one plausible pronunciation based on orthographic bodies in English (i.e., -ink as in “sink”). MM pseudowords (e.g., “chead”) involve at least two plausible pronunciations based on existing orthographic bodies in English (i.e., -ead pronounced either as in “bead” or “head”). Participants were instructed to read each pseudoword as quickly and accurately as possible. Pseudowords were displayed in blocks of 20 items, with self-paced breaks between blocks. If the participant did not respond within 10 seconds, the trial timed out, and the participant was prompted to proceed to the next item. All stimuli were monosyllabic and 3-6 letters in length. Accuracy was scored for the first complete attempt. Any plausible pronunciation of each pseudoword based on English spelling-sound correspondences was scored as correct.

#### Semantics composite score

The stroke survivors completed a battery of tasks designed to assess the intactness of their semantic representations and processing. Three tasks assessed semantic processing without requiring reading or verbal responses: Pyramids and Palm Trees (Howard & Patterson, 1992), the Temple Assessment of Language and Short-term Memory in Aphasia (TALSA) category judgment subtest (Martin, Minkina, Kohen, & Kalinyak-Fliszar, 2018), and an auditory word-to-picture matching task. For Pyramids and Palm Trees, the subject was presented a picture on a touchscreen (e.g., a pyramid) and matched it to one of two pictures presented below: the semantically related target (e.g., a palm tree) or the incorrect distractor (e.g., a pine tree; 30 trials). The TALSA category judgment subtest measured knowledge of animals, transportation, vegetables, furniture, and musical instruments through 60 pairs of pictures. Two pictures were displayed in succession for three seconds each (e.g., banana and apple), and the subject had to indicate whether the two pictures belonged to the same category by pressing “yes” or “no” on the touchscreen. For auditory word-to-picture matching, the participant was presented with a word auditorily via headphones (48 trials) and selected the corresponding picture among five semantically related foils on a touchscreen.

Two picture naming tasks assessed stroke survivors’ ability to generate phonology from semantics: the 60-item Philadelphia Naming Test (Roach et al., 1996) and an in-house 60-item picture naming test (Fama et al., 2019). Participants were instructed to use one word to name aloud each picture as quickly and accurately as possible, within a 20-second limit. Accuracy was scored on the first complete attempt.

Accuracy on each of these five tasks was averaged to produce a semantic composite score for each person. Three participants were missing PNT scores and one participant was missing a PPT score, so their composite scores reflected four tasks. One participant was missing both an auditory word-to-picture matching score and a PNT score, so their composite score reflects three tasks.

#### Phonology composite score

We also collapsed across five tasks that assessed phonological processing to obtain a phonology composite score for the stroke survivors. These tasks assessed the stroke survivors’ capacity to process and manipulate speech sounds. Three tasks required no verbal output. The TALSA phoneme discrimination task required participants to distinguish between pairs of pseudowords that were either the same or differed by 1-3 syllables (20 trials) (Martin et al., 2018). Two rhyme judgement tasks (auditory, picture) assessed the ability to access phonological representations (Fama et al., 2019). In the auditory version, participants heard pairs of prerecorded words (40 trials) and pressed “yes” or “no” on the touchscreen to indicate if they rhymed. In the picture rhyme judgement task, participants were presented with pairs of images (40 trials) and used the touchscreen to indicate if they rhymed.

Two tasks required the participants to provide verbal responses: a pseudoword repetition task and a rhyme production task. In the pseudoword repetition task, participants heard an auditorily-presented pseudoword and had to repeat it aloud (60 trials). In the rhyme production task, stroke survivors were auditorily presented with a word and were required to produce a rhyming word (25 trials).

Again, accuracy on the five tasks were averaged to produce a phonology composite score. Two participants were missing pseudoword repetition scores and one participant was missing a rhyme production score, so their composite scores reflected four tasks.

#### Motor speech assessment

Stroke survivors were also assessed for motor speech deficits using the Apraxia of Speech Ratings Scale 3.0 (ASRS3). This assessment was critical, as the model is not designed to accommodate motor speech deficits, and their presence could bias how well the models can simulate alexia. Stroke survivors were given a severity rating from 0 (indicating “not present) to 4 (“nearly always evident”) (Strand et al., 2014; Wambaugh et al., 2019). Ratings were done via video review by a certified speech-language pathologist. For stroke survivors with motor speech impairments, their oral responses on all tasks were scored with a motor speech leniency. They were given credit if their response included a prolongation, segmentation, or distortion that did not cross phoneme boundaries. Motor speech leniency was not given for additions (i.e., intrusive schwa due to apraxia), substitutions, or distortions that impacted the intelligibility and thus transcription of the utterance.

### Neuroimaging

#### MRI acquisition and lesion tracing

MRIs in this study are used only to determine lesion volume for statistical analyses. The following sequences were acquired with Georgetown’s 3T Siemens MAGNETOM Prisma scanner using a 20-channel head coil, as previously described in Dickens et al. (2021): a T1-weighted magnetization prepared rapid gradient echo (MPRAGE) sequence (1 mm^3^ voxels), a fluid-attenuated inversion recovery (FLAIR) sequence (1 mm^3^ voxels), and a high angular resolution diffusion imaging (HARDI) sequence (81 directions at *b* = 3000, 40 at *b* = 1200, 7 at *b* = 0; 2 mm^3^ voxels). For stroke survivors, a board-certified neurologist (author P.E.T.) manually traced stroke lesions on the native-space MPRAGE and FLAIR images using ITK-SNAP (Yushkevich et al., 2006; http://www.itksnap.org/). MPRAGEs and lesion tracings were warped to the Clinical Toolbox Older Adult Template (Rorden, Bonilha, Fridriksson, Bender, & Karnath, 2012) using the Advanced Normalization Tools toolbox (Avants et al., 2011) as described in Dickens et al. (2019). Further details can be found in Dickens et al. (2021). Lesion volume was calculated as the count of voxels in an individual’s lesion map in standardized brain template space. A lesion overlap image illustrates that lesions in the sample (Fig. 1).

**Fig. 1.**
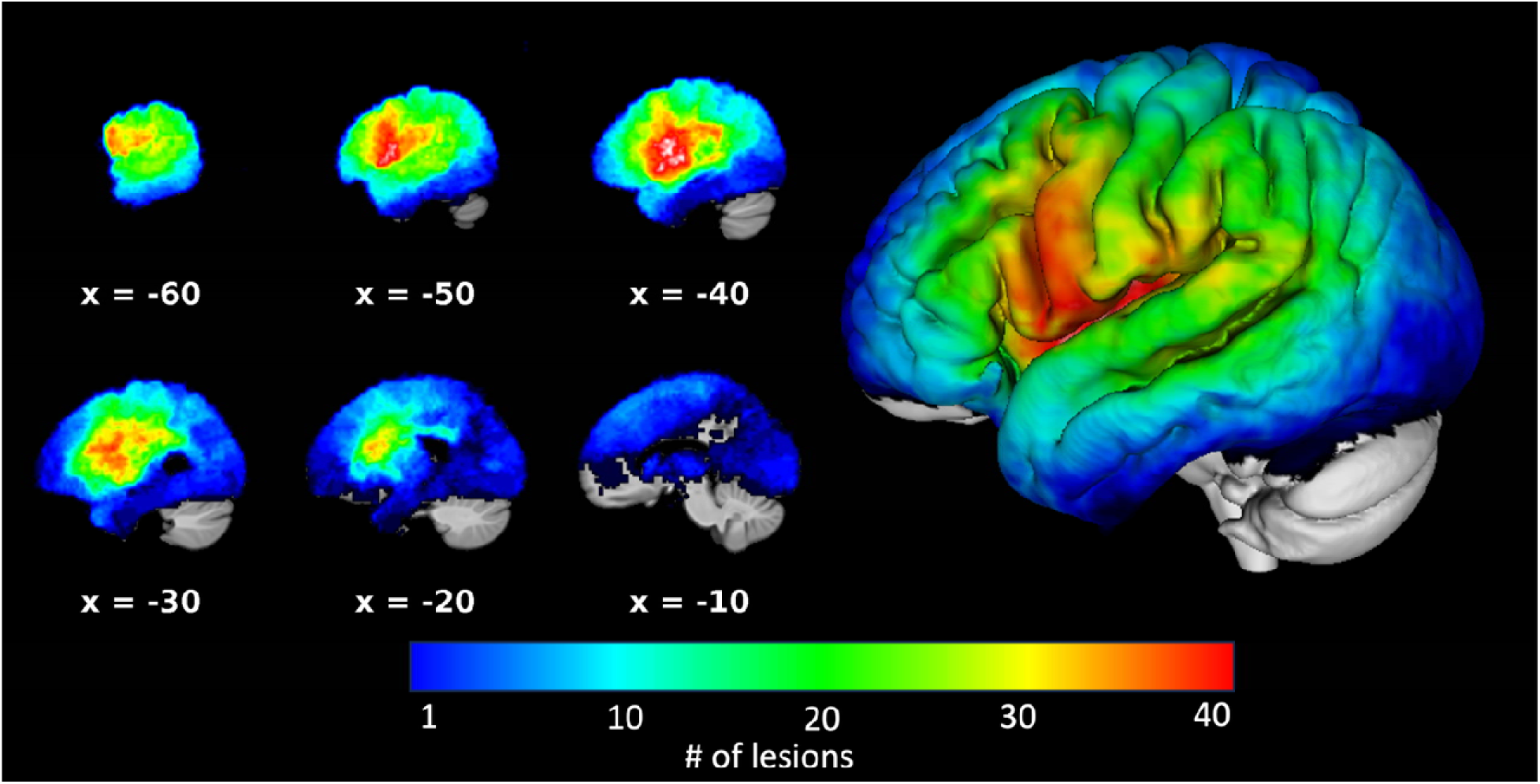
– Lesion overlap map. The lesion overlap map demonstrates that lesions were concentrated in perisylvian regions involved in language and reading.

### Statistical Analyses

#### Oral Reading

To determine differences between reading scores of controls and stroke survivors, we analyzed the oral reading data using two logistic mixed-effects models. The first model assessed main effects of patient status, frequency, and consistency, as well as the two- and three-way interaction of those factors, on reading accuracy. Age and education were included as covariates of no interest with random intercepts for subject and item. The second model assessed the main effects of patient status and lexicality on reading accuracy, including fixed effects of age and education as covariates and random intercepts of subject and item. Models were fit using the *glmer* function of the lme4 package (v1.1-35.4) in R Statistical Software (v4.4.1) using maximum likelihood estimation. Model results are reported as odds ratios with 95% confidence intervals. Statistical significance was determined by Wald Z-statistics and their associated p-values.

#### Semantic and phonological composite scores

To assess if the stroke survivors have isolated or co-occurring phonological and semantic impairments, we calculated the correlation of their phonological and semantic composite scores.

### Computational Modeling

We implemented the triangle model of single word reading from Harm & Seidenberg (2004) using recurrent artificial neural networks, with one modification (see below). These models develop “hidden” representations that map between prespecified inputs and outputs via a learning algorithm – here, recurrent backpropagation through time over five intervals of time.

#### Model Architecture

The model (Fig. 2) has two pathways for reading words: a direct, orthography-phonology pathway reflecting the sublexical reading pathway, and an indirect, orthography-semantics-phonology pathway, reflecting the lexical reading pathway. The 105-unit orthography input layer projects to a 61-unit phonology output layer via a 100-unit orthography-phonology (OP) hidden layer, and to a 300-unit semantic output layer via a 500-unit orthography-semantics (OS) hidden layer. The semantic layer connects to the phonology layer via a 100-unit semantic-phonology (SP) hidden layer, and the reverse phonology-semantics connection is implemented by a separate 500-unit phonology-semantic (PS) hidden layer. Finally, both the phonology and semantic output layers have recurrent “Elman” connections, which allow the previous time interval (activation of the pertinent layer at time *t*-1) to influence activation at time *t*. The Elman layers are the only architectural difference between the present model and that in Harm & Seidenberg (2004), replacing the continuous “clean up” layers in their model. Both the continuous layers used in Harm & Seidenberg (2004) and the Elman layers used here serve the same purpose: they pull a layers’ inputs into a smaller number of learned “attractors”, which are learned over training and constitute correct output states. Attractors can thus both assist in learning and also allow for variations in input to be mapped onto an invariant output. Elman layers, rather than the continuous units used in Harm & Seidenberg (2004), were implemented because they simplify the time dynamics of the model, which are not critical to the goal of the present research, and thus reduce computational demands.

**Fig. 2.**
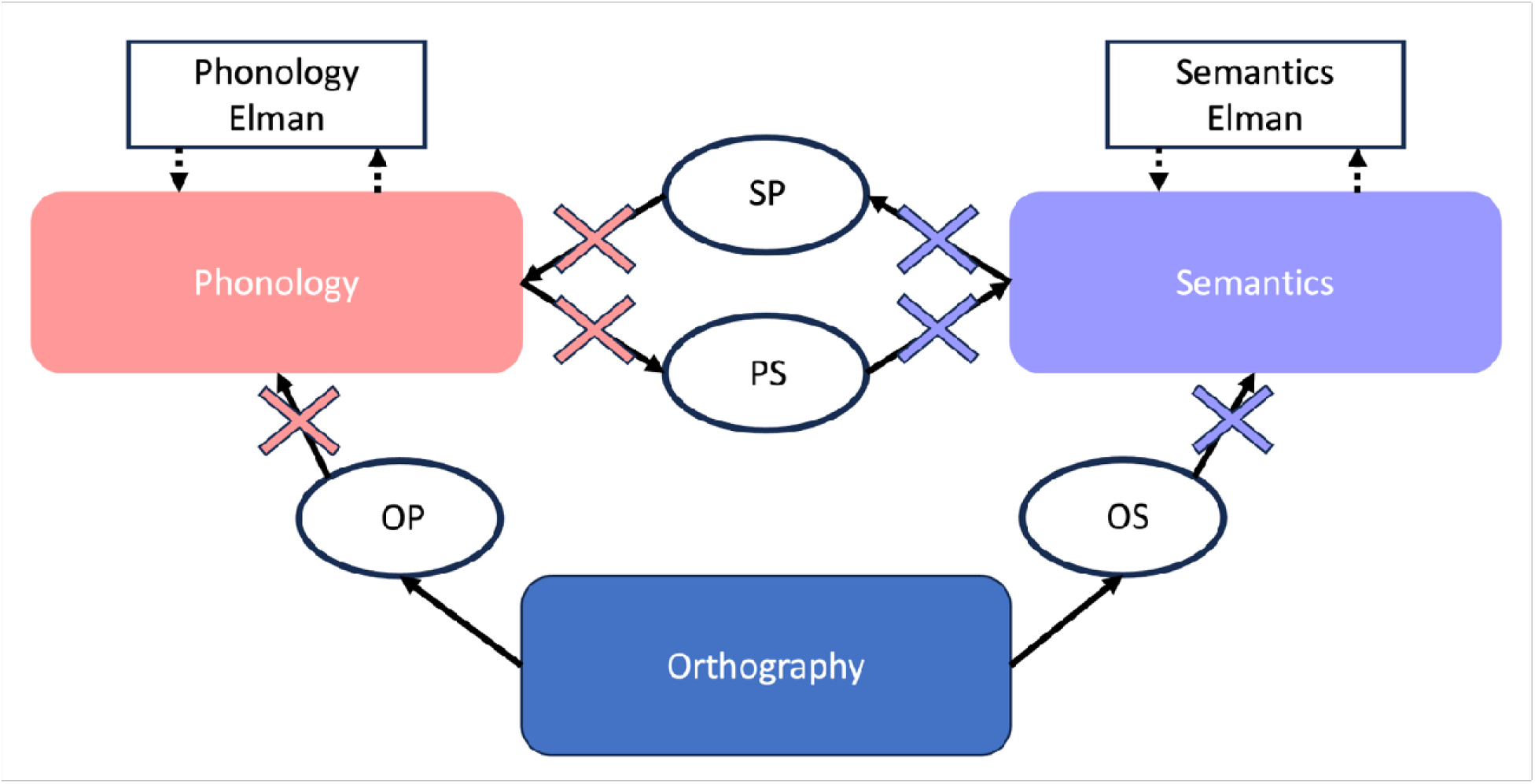
– Model diagram. Rounded rectangles indicate input and output layers, which have predetermined representations. Ovals indicate hidden layers, which develop intermediate representations over the course of training. Rectangles indicate Elman layers, which implement recurrent processes in the model by feeding the activations of the connected layers at time *t*-1 as input on time *t*. Light red ×s indicate location of links removed to implement phonological lesions. Light blue ×s indicate location of links removed to implement semantic lesions.

#### Model Corpora

The corpus used to train the model (the Training Corpus) was the 2998-word corpus from Plaut et al., (Plaut et al., 1996). Briefly, the Training Corpus covers most of the single-syllable words in English. Contained within the Training Corpus are four subsets of words used to evaluate the model’s accuracy on frequency and consistency, taken from Taraban & McClelland, (1987). The two consistent Real Word Test sets (high frequency consistent, low frequency consistent) have 24 items each. The two inconsistent Real Word Test sets (high frequency inconsistent, low frequency inconsistent) have 48 items each. The inconsistent Real Word Test sets have more words than the consistent Real Word Test Sets because both Taraban & McClelland (1987) and Plaut et al. (1996) were interested in cases of “regular inconsistent” words, whose pronunciation is similar to many but not all orthographic neighbors, where we did not distinguish between regular inconsistent and irregular words.

The model was also tested on its ability to read single-syllable pseudowords. The Pseudoword Test set consisted of 166 items, compiled from Glushko (1979) and Taraban & McClelland (1987).

#### Model Representations

Orthographic and phonological word representations were the same as those used in Plaut et al., (1996). Briefly, the representations were binary, slot-based grapheme and phoneme representations, divided into onset, vowel and coda. The semantic features were generated using the COALS algorithm (Rohde, Gonnerman, & Plaut, 2006) from the Google Web 1T 5-gram Version 1 dataset. In short, word co-occurrences within a 5-word ramped window are converted into pairwise correlations. We use semantic representations incorporating a modification based on Chang et al., (2012), where a 150-unit base vector is generated and then duplicated to produce a 300-unit representation. The largest positive *n* values in the first half and the largest *m* negative values in the second half are set to 1, and all others are set to 0. Following Chang et al., (2012), we used *n* = 3 and *m* = 7, producing representations that uniquely specified all words in the Training Corpus.

#### Model Training

The models were trained in three phases: Preliterate, Reading, and Recovery. In all phases, training was online (weights were updated after each stimulus presentation), with stimuli selected from the Training Corpus for a given training instance with a probability relative to their logarithmically-compressed frequency (Plaut et al., 1996). Across all three phases, every instance of training was identical. First, an item was probabilistically selected from the Training Corpus. The representation for that item was “clamped” onto the appropriate input layer: the activations of the input layer were set to the pattern of ones and zeros that specify the representation, and not allowed to change. The model cycled for five time steps. At each time step, unit activations were updated. After the last time step, the binary cross entropy between the model’s activation at the relevant output layer(s) and the correct representation of the word being trained was measured. Weights were then updated via backpropagation through time, with the magnitude of the weight change being proportional to the measured binary cross entropy.

The Preliterate phase simulates the development of expressive and receptive speech processing, which occurs before children learn to read. In the Preliterate phase, the models were trained on comprehension and speech tasks. In the comprehension task, phonological input was clamped onto the phonological layer, and the model was trained to produce the correct semantics. Only the connections from phonology to semantics were trained. The speech task was the inverse of the phonology task: semantic input was clamped onto the semantic layer, and the model was required to produce the correct phonology. Only the connections from semantics to phonology were trained. During the Preliterate phase, the links from orthography to semantics and to phonology were present, but the weights were not updated. The preliterate training phase lasted for 1.5 million examples (750,000 examples for comprehension and 750,000 for speech).

After the Preliterate training phase, the phonology-to-semantics and semantics-to-phonology weights were frozen and the models proceeded to the Reading phase. Orthographic inputs were clamped onto the orthography layer, and the model was required to generate correct phonological and semantic outputs. Only the orthography-phonology and orthography-semantics connections were trained during this phase. The Reading phase lasted for 750,000 examples^1^.

The Recovery phase, which occurred after the models were lesioned (see below), was identical to the Reading phase except in the number of examples. In the Recovery Phase, the models were trained for 50,000 examples.

All units in the model used sigmoid activation functions. A learning rate of .05 and weight decay of 1E-7 were used throughout. A constant, untrainable bias of -2 was applied to all layers to enforce sparse activations and to prevent early, strong activations from unduly influencing training (Ueno, Saito, Rogers, & Lambon Ralph, 2011). In the Preliterate phase, no error was backpropagated from an output unit (and thus no weights were modified) if its activation was within .1 of the target. In the Reading phase, no error was backpropagated when unit activation was within .001 of the target (Chang, Welbourne, Furber, & Lambon Ralph, 2024). This “zero error radius” helps prevent overfitting. To avoid any idiosyncratic model from unduly influencing the results, five independent healthy models were trained. Their results were averaged to assess “healthy” reading in the model, but the models were separately lesioned for the purpose of simulating alexia (see below).

#### Model Lesioning

The trained models were lesioned in their capacity to process phonology, semantics, or both. Lesions were accomplished by removing percentages of the incoming and outgoing connections to the phonology and/or semantic layers. For example, a 50% lesion to phonology consisted of the instantaneous removal of 50% of the links connecting the OP hidden layer to phonology, 50% of the links connecting the SP layer to phonology, and 50% of the links connecting the phonology layer to the PS layer (light red ×s in Fig. 1). Nineteen levels of damage (0% to 95%, in 5% intervals) were applied to the semantic and phonological layers separately. All possible combinations of lesions to the phonology and semantic layers were applied, resulting in 400 unique lesion combinations. These 400 lesions configurations were applied to each of the 5 trained models. Each of the five healthy models were lesioned 3 times at each level of damage (i.e. 3 * 400). Each specific instance of the 15 models at each lesion configuration used a unique seed to select the random subset of connections that were removed. The results of the 15 damaged models at each lesion combination were averaged to produce the behavior for that particular lesion configuration. After lesioning, each model was retrained for 50,000 presentations to allow for plasticity-related recovery (Welbourne & Lambon Ralph, 2005).

#### Model Testing

After each phase of training, the model’s activations at the relevant output layer(s) for every word in the Training Corpus were recorded at the last time interval. Outputs were binarized such that activations at or above .5 were set to 1 and activations below .5 were set to 0 (Plaut et al., 1996). Binarizing the raw model outputs allows for unambiguous interpretation of which phonemes or semantic features are activated in response to a given input: either a unit is on or off.

The models’ oral reading accuracy was assessed using the outputs of the phonology layer. The binarized phonological output was assessed as in Plaut et al., (1996). Onset and codas were divided into mutually exclusive slots, and the most active unit activated to .5 or higher was selected as that slot’s output. When no unit was activated at or above .5, that slot had no output. For the vowel slot, the most active vowel unit, regardless of level of activation, was selected as the output. For word examples, the model output was assessed as correct only if it exactly matched the appropriate phonological or semantic representation. For pseudoword examples, the phonological output was assessed as correct if it matched any grapheme-phoneme correspondences that were found in the Training Corpus. This matches the way the pseudoword reading test was scored for stroke survivors, in which any pronunciation that was plausible based on English spelling-sound correspondences was scored as correct.

We also assessed if the model generated the correct semantic outputs for each word. Because semantic access from text was not the focus of the current research, we used a permissive measure of accuracy. Semantic outputs were binarized in the same way as the phonological outputs. Semantic accuracy was then assessed by computing the Euclidean distance between the generated output and every word representation in the Training Corpus (Chang, 2023; Monaghan, Chang, Welbourne, & Brysbaert, 2017). If the smallest obtained Euclidean distance was between the generated output and the correct word representation, the output was marked correct.

#### Confirming that healthy models matched healthy reading

To verify that the healthy models provided a good fit to the pattern of reading ability shown by the healthy controls, the reading accuracies within each word type (all real words, pseudowords, and the Real Word Test sets) were submitted to two-sample t-tests.

#### Assessing Psycholinguistic Effects in the Models

To determine if the lesioned models produced the expected psycholinguistic effects, we analyzed the full set of 400 lesioned models with two linear mixed effects models. The first model examined frequency and consistency effects, predicting accuracy from the main effects and interaction of frequency and consistency, with a random intercept of model number. The second model examined the lexicality effect, predicting accuracy from word type (real word or pseudoword) with a random intercept of model.

#### Assessing the Effects of Semantic and Phonological Lesions on Reading in the Models

To assess the effects of semantic and phonological damage on reading accuracy in the models, we conducted two linear mixed effects models. The first predicted reading accuracy from fixed main effects of frequency, consistency, semantic lesion severity, and phonological lesion severity, as well as the two-, three-, and four-way interactions between these effects. The intercept of model number was included as a random effect.

The second linear mixed effects model predicted accuracy from the fixed main effects of lexicality (word, pseudoword), semantic lesion severity, and phonological lesion severity, as well as the two- and three-way interactions between these effects. The intercept of model number was included as a random effect.

#### Matching Models to Individual Stroke Survivors

To match a given lesioned model with a specific stroke survivor, the Euclidean distance was computed between each stroke survivors’ two-feature vector of word and pseudoword reading accuracy and the same vector for the 400 lesion configurations. We chose to match the models to stroke survivors based on word and pseudoword reading because they represent the most robust and specific measures of semantic versus phonological reading. The model producing the smallest Euclidean distance was selected as the “matched model” for that stroke survivor, meaning that the semantic and phonological lesion severities producing the most similar behavior were taken as representative of that individual’s impairments to the cognitive processes underlying reading aloud. We take the inverse of the matched-model lesion severities (1 - % lesioned links) as the model’s phonological and semantic lesion scores. This preserves an intuitive relationship such that larger model lesion scores (indicating more intact processing capacity in the model) are analogous to larger composite scores (indicating more intact cognitive processes in the stroke survivors).

#### Assessing Behavioral Matches

We assessed the stroke survivor-model matches by computing separate correlations between the stroke survivor and matched-model reading of each of the Real Word Test sets (high and low frequency/consistency words), across the sample. These correlations assessed how well the models captured reading of each word type, across the sample. Because the representations in the model were not designed to simulate imageability, we collapse across high and low imageability throughout. Effects related to letter length and articulatory complexity are not critical to the assessment of alexia, so we also did not attempt to simulate them.

Identifying and understanding systematic differences between the real and simulated scores permit understanding where the models fail to capture reading, potentially leading to a greater understanding of reading as a cognitive process and targets for improving the models. To assess how well the matched models simulated the numerical values of stroke survivors’ accuracy, we calculated the difference (stroke survivors’ reading accuracy – matched models’ reading accuracy) within each word type (all words, pseudowords, and the four Real Word Test Sets). The computed differences were submitted to a paired t-test. We also computed the absolute value of the differences within each word type, allowing us to assess the total error. The absolute differences were submitted to a one-sample t-test against zero.

We also computed the correlation between the model lesion scores and the directly-measured phonological and semantic composite scores. Because we did not directly simulate semantic and phonological tasks, the direction of relationship between the model lesion scores and the directly-measured composite scores is pertinent. However, there is no reason to expect the scores to be on the same scale.

#### Assessing the sources of differences between stroke survivors’ and matched-model reading accuracy

We aimed to identify if poor fits between the stroke survivors and matched-models are due to demographic or cognitive variables. To do so, we estimated a linear mixed effects model predicting mean deviance from the main effects, as well as two- and three-way interactions of word type (categorical: high/low frequency and consistency), semantic composite score, and phonological composite score. Age, education in years, and chronicity in months were included as demographic predictors. An intercept of subject was included as a random effect. The ASRS3 Apraxia of Speech Severity score was included as a covariate of no interest. All continuous variables were mean-centered to enable interpretation of word type effects via estimated marginal means. The significance of fixed effects was evaluated via Wald tests. We tested the conditional marginal mean by word type against zero, the value at which a model would perfectly match a stroke survivor’s reading accuracy. To evaluate if semantic and phonological composite score affected deviance, we tested the marginal mean of deviance by word type at +/-1 SD for both scores. All tests were FDR corrected for multiple comparisons.

## Results

### Profiles of reading and cognitive deficits in the stroke group

We first confirmed if the stroke survivors exhibited the standard psycholinguistic effects of frequency and consistency. Stroke survivors read aloud all word types less accurately than matched controls (Table 2, Fig. 3). A logistic mixed-effects model showed expected main effects of group (Z = -8.92, p < .001, OR = .010, 95% CI = 0.00-0.03), frequency (Z = -3.39, p = .001, OR = 0.20, 95% CI = 0.08-0.51), and consistency (Z = -4.76, p < .001, OR = 0.12, 95% CI = 0.05-0.28), such that controls read more accurately than stroke survivors and higher frequency words and consistent words were read more accurately across both groups. There was also an effect of age (Z = 2.51, p = .012, OR = 1.04, 95% CI = 1.01-1.06), such that older participants were more accurate. Additionally, there was an interaction of group and consistency (Z = 4.19, p < .001, OR = 5.61, 95% CI = 2.50-12.58), such that stroke survivors were less accurate relative to controls at reading inconsistent words (Supplementary Table 1; Supplementary Fig. 1). Notably, there was no interaction of group by frequency by consistency, showing that these participants do not show the disproportionately poor performance on low-frequency inconsistent words canonically expected in surface alexia.

**Fig. 3.**
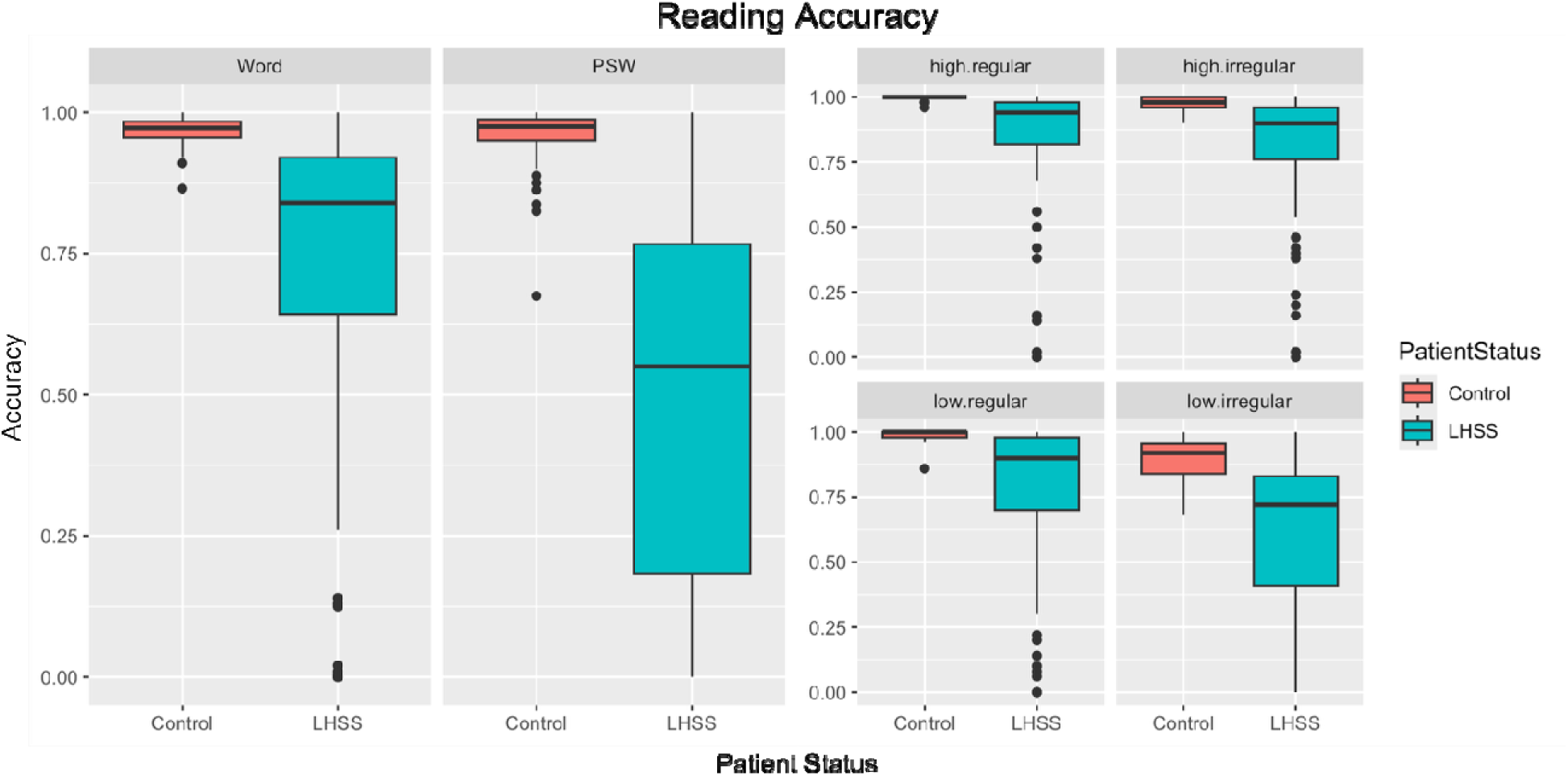
– Reading accuracies of controls and stroke survivors. PSW = pseudoword, LHSS = left-hemisphere stroke survivor.

**Table 2.**
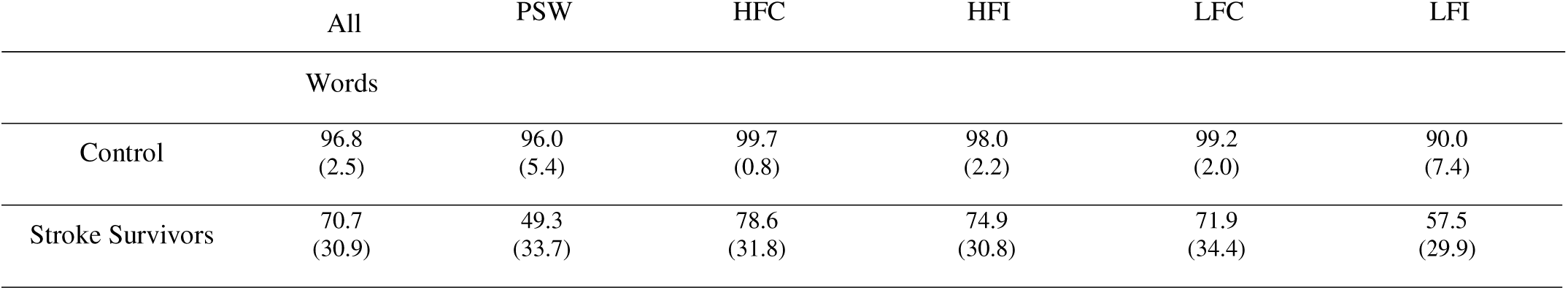
– Control and stroke survivor reading accuracies.

We then confirmed that stroke survivors showed the standard effect of lexicality. A second logistic mixed-effects model examining the effect of group and lexicality on oral reading accuracy revealed expected main effects of group (Z = -11.86, p < .001, OR = 0.02, 95% CI = 0.01-0.03) and lexicality (Z = 4.59, p < .001, OR = 2.28, 95% CI = 1.60-3.25), such that controls read more accurately than stroke survivors and real words were read more accurately than pseudowords across both groups (Supplementary Table 2; Supplementary Fig. 2). This effect was confirmed by a significant interaction of group by lexicality (Z = 9.17, p < .001, OR = 2.72, 95% CI = 2.19-3.36). This exaggerated lexicality effect is typical of phonological alexia.

We next assessed semantic and cognitive processing in the stroke survivors to understand if their reading deficits reflect isolated (i.e., just semantic or phonological) or co-occurring cognitive deficits. Consistent with the expected cognitive outcomes of left hemisphere stroke, stroke survivors were broadly more phonologically impaired than semantically impaired (Table 3). However, phonology and semantic composite scores were highly correlated (*r* = 0.76, *p* < .001, Fig. 4a). To determine if this was an effect of larger lesions or demographic variables, we separately residualized the phonology and semantic composite scores on lesion volume, age, education, and ASRS 3.0 Apraxia of Speech (AoS) severity score. Accounting for this variance decreased this correlation, but it remained strong (*r* = 0.67, *p* < .001, Supplementary Fig. 3), showing that these participants do not show the isolated lesion patterns that might be expected to underlie pure phonological or surface alexia. Instead, they show a mixed presentation. Despite this correlation, substantial variation in the degree of phonological vs. semantic impairments between individuals still exists (Fig 4a).

**Fig. 4.**
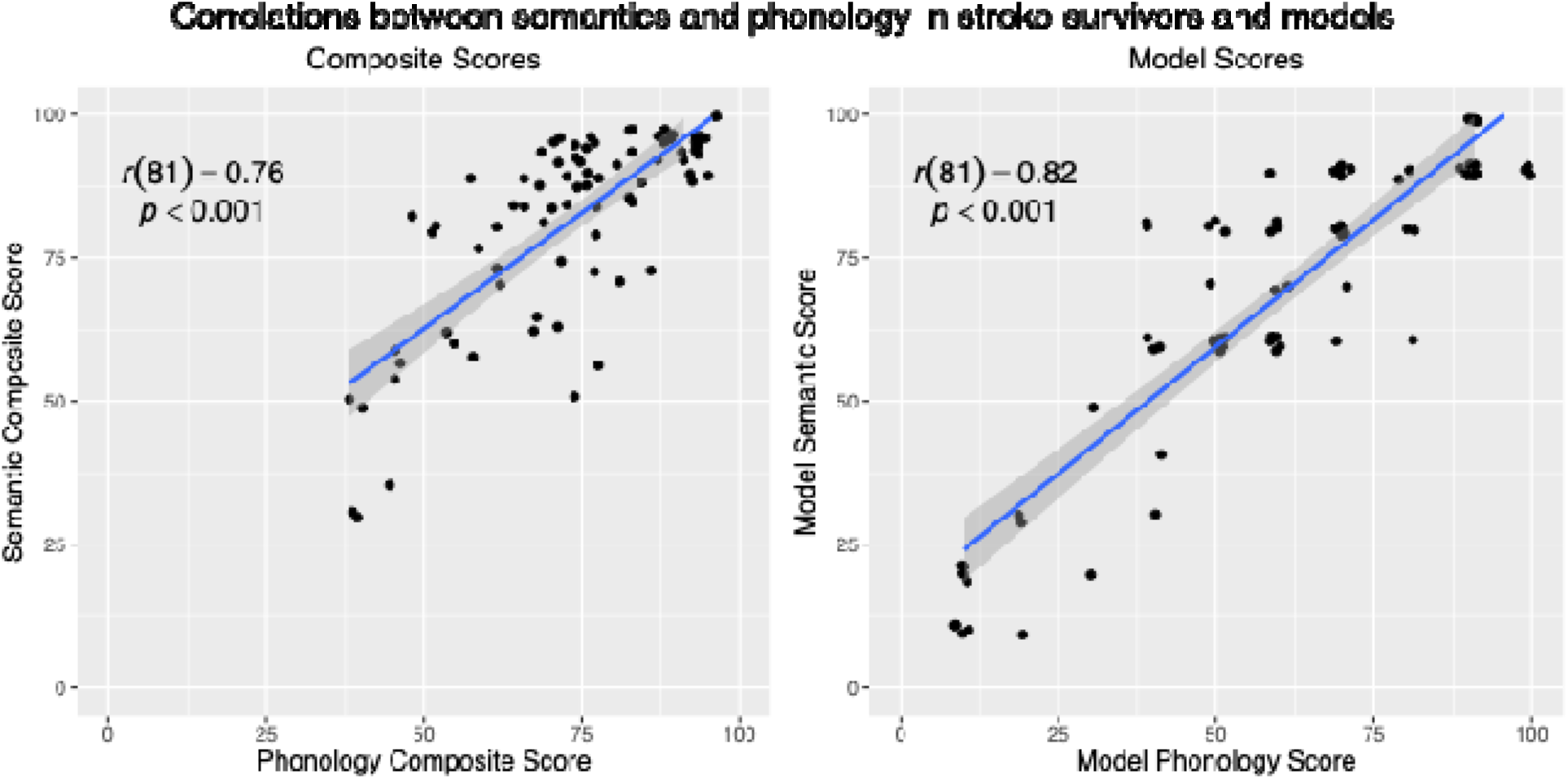
– Correlations between the semantics and phonology composite scores (Panel A) and the model semantics and phonology scores (Panel B). A small amount of jitter has been added to Panel B to accommodate clusters of identical model lesions, which occurred due to the discrete nature of the model lesions.

**Table 3.**
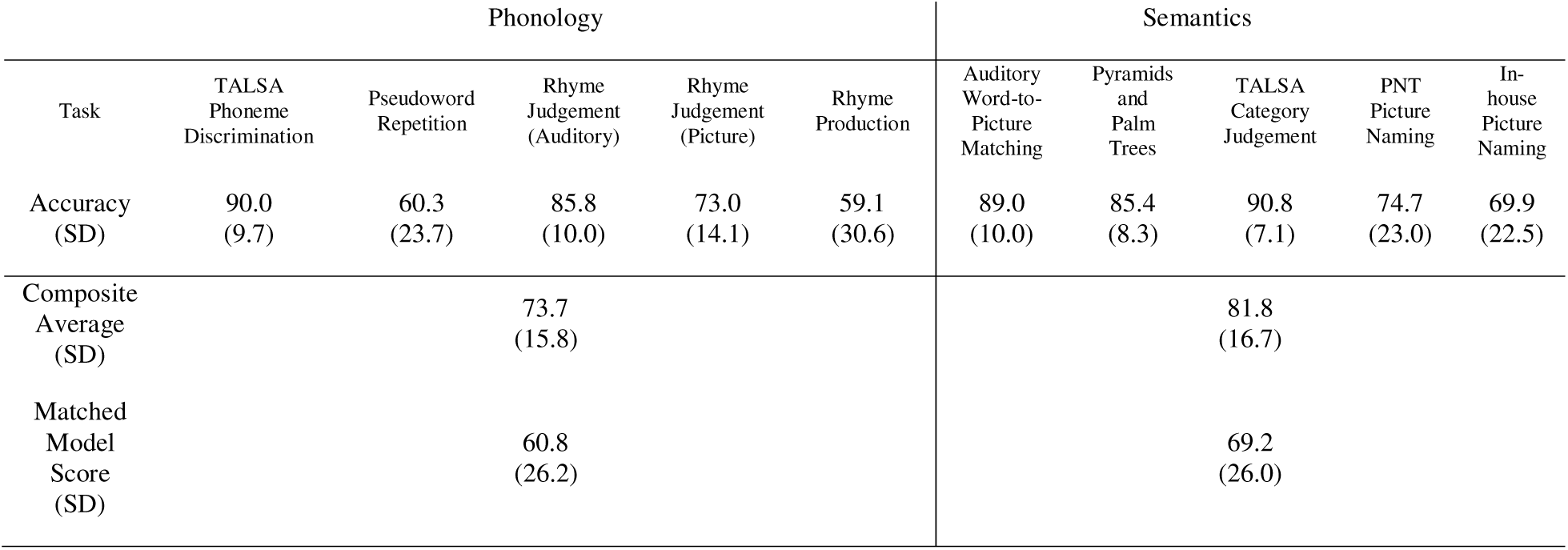
– Performance on semantic and phonological tasks.

| Task | Phonology |  |  |  |  | Semantics |  |  |  |  |
| --- | --- | --- | --- | --- | --- | --- | --- | --- | --- | --- |
|  | TALSA<br>Phoneme<br>Discrimination | Pseudoword<br>Repetition | Rhyme<br>Judgement<br>(Auditory) | Rhyme<br>Judgement<br>(Picture) | Rhyme<br>Production | Auditory<br>Word-to-Picture<br>Matching | Pyramids<br>and<br>Palm<br>Trees | TALSA<br>Category<br>Judgement | PNT<br>Picture<br>Naming | In-house<br>Picture<br>Naming |
| Accuracy<br>(SD) | 90.0<br>(9.7) | 60.3<br>(23.7) | 85.8<br>(10.0) | 73.0<br>(14.1) | 59.1<br>(30.6) | 89.0<br>(10.0) | 85.4<br>(8.3) | 90.8<br>(7.1) | 74.7<br>(23.0) | 69.9<br>(22.5) |
| Composite<br>Average<br>(SD) | 73.7<br>(15.8) |  |  |  |  | 81.8<br>(16.7) |  |  |  |  |
| Matched<br>Model<br>Score<br>(SD) | 60.8<br>(26.2) |  |  |  |  | 69.2<br>(26.0) |  |  |  |  |

### Confirming the model simulates reading in control participants

The unlesioned models replicated the qualitative pattern of accuracy across word types in our sample of healthy controls (Table 4). Reading accuracy overall, as well as HFI, LFC, and LFI reading did not differ between the healthy controls and unlesioned models (*p*’s < .05). However, the models did perform worse on pseudowords (*t*(5.1) = -4.5, *p* = .006) and slightly better on high frequency consistent words (*t*(60) = 2.6, *p* = .010) than the controls did. The difference in HFC word accuracy was because none of the models made any errors to words in the test set. While the models read pseudowords less accurately than the controls, the obtained accuracy was in line with previous neural network models of reading (Chang et al., 2024; Harm & Seidenberg, 2004; Plaut et al., 1996), and captured the pattern of less accurate pseudoword than real word reading.

**Table 4.** – Control participant and unlesioned model reading accuracies.

|  | All Words | PSW | HFC | HFI | LFC | LFI |
| --- | --- | --- | --- | --- | --- | --- |
| Control | 96.8<br>(2.5) | 96.0<br>(5.4) | 99.7<br>(0.8) | 98.0<br>(2.2) | 99.2<br>(2.0) | 90.0<br>(7.4) |
| Healthy Model | 97.6<br>(0.6) | 88.4<br>(3.3) | 100.0<br>(0.0) | 94.8<br>(4.0) | 97.9<br>(2.9) | 88.5<br>(4.0) |
| $t$<br>(d.f.) | 1.71<br>(18.6) | -4.5<br>(5.1) | 2.65<br>(60) | -1.9<br>(4.3) | -1.5<br>(4.3) | -1.2<br>(6.3) |
| $p$ | 0.103 | 0.006 | 0.010 | 0.124 | 0.208 | 0.27 |

PSW = pseudowords, HFC = high frequency consistent words, HFI = high frequency inconsistent words, LFC = low frequency consistent words, LFI = low frequency inconsistent words

### Psycholinguistic effects in the lesioned models

We estimated two linear mixed effects models, one to confirm that the lesioned models produced the expected frequency by consistency interaction and the second to assess the lexicality effect on accuracy. The frequency by consistency model showed main effects of both frequency (β = -0.03, *t*(294) = -4.37, *p* < .001) and consistency (β = 0.07, *t*(294) = 11.95, *p* < .001), as well as the interaction of frequency and consistency (β = 0.02, *t*(294) = 2.22, *p* = .027). High frequency words and consistent words were read more accurately than low frequency or consistent words, and low frequency inconsistent words were read less accurately than any other combination of factors (Supplementary Table 3; Supplementary Fig 4). A second model examined the effect of lexicality. Accuracy was predicted by word type (β = 0.12, *t*(98) = 15.41, *p* < .001), such that real words were read more accurately than pseudowords (Supplementary Table 4; Supplementary Fig. 5).

### Effects of model lesions on model reading performance

Two linear mixed effects models estimated the impact of semantic and phonological lesions on model reading. The first model assessed how these lesions impacted reading real words of high or low frequency and consistency (Supplementary Table 5; Supplementary Fig. 6. There were main effects of consistency (β = -0.04, *t*(285) = -2.32, *p* = .021), semantic lesion severity (β = -0.42, *t*(125) = -8.45, *p* < .001), and phonological lesion severity (β = -1.26, *t*(125) = -25.13, *p* < .001). Inconsistent words were read less accurately, and larger semantic and phonological lesions produced less accurate reading. Three two-way interactions were obtained. There was an interaction of consistency and phonological lesion severity (β = 0.06, *t*(285) = 2.13, *p* < .034), consistency and semantic lesion severity (β = 0.24, *t*(285) = 8.23, *p* < .001), and semantic and phonological lesion severity (β = 0.26, *t*(285) = 2.79, *p* = .006). Larger semantic lesions caused worse inconsistent word reading. Larger phonological lesions caused worse consistent word reading relative to inconsistent word reading. Finally, smaller semantic lesions caused a smaller decrease in reading accuracy when phonological lesions were small, relative to when phonological lesions were large. Finally, there was a three-way interaction of consistency, semantic lesion severity, and phonological lesion severity (β = -0.14, *t*(284) = -2.53, *p* < .012). Semantic lesions decreased reading accuracy on inconsistent words to a greater degree when phonology lesions were large, relative to when they were small. In sum, semantic lesions had a disproportionate effect on inconsistent words, where phonological lesions had a disproportionate effect on consistent words (Fig. 5). The main effect of frequency disappeared in this model, and frequency did not interact with any other predictors. This indicates that the frequency effect is weaker in the models than it is in the stroke survivors.

**Fig. 5.**
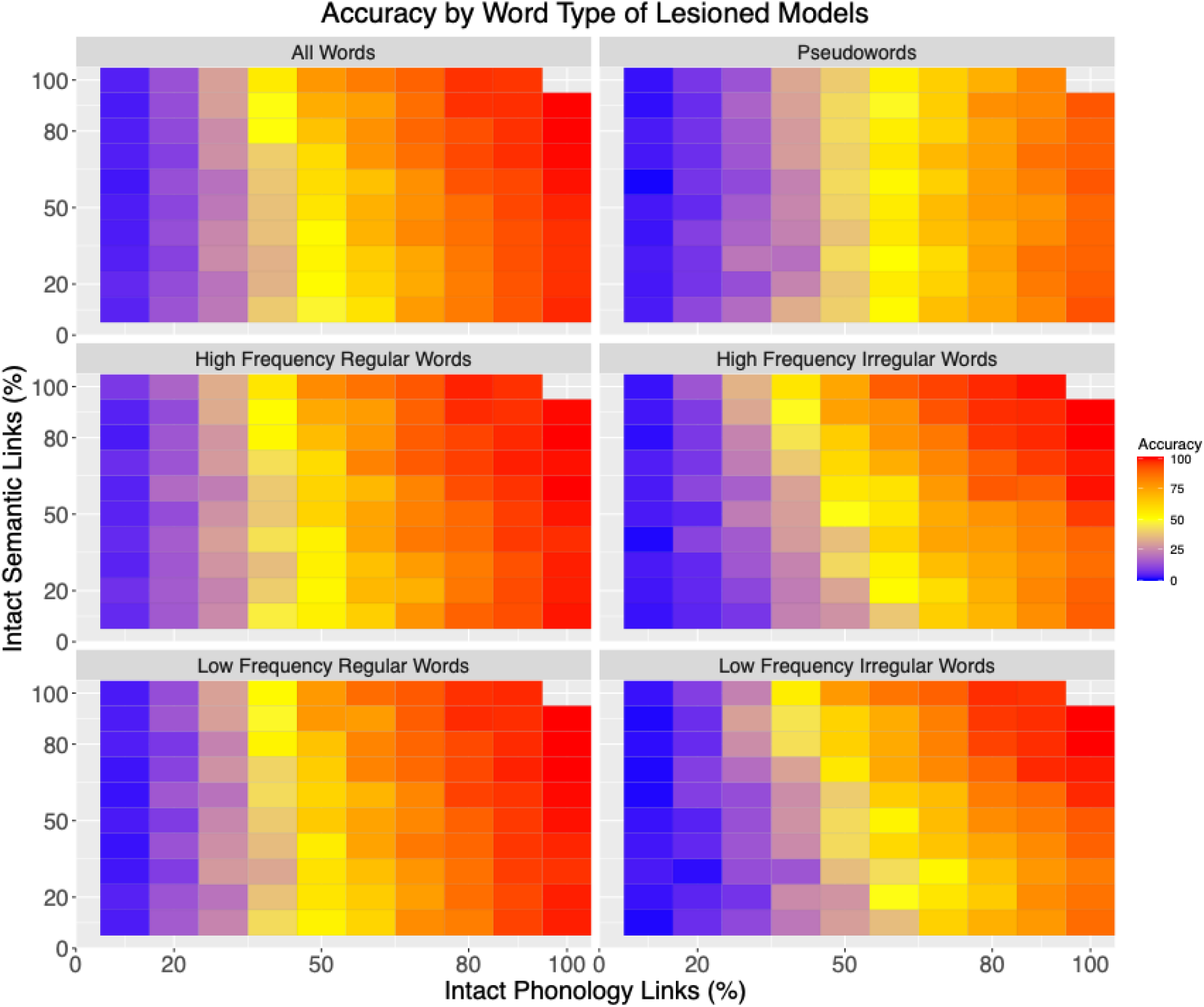
– Reading accuracy of lesioned models by word type. Figures show model reading accuracy for each word type as a function of lesion load on phonology and semantics. Real words display a diagonal pattern, showing that both semantic and phonological lesions affect accuracy, most apparent with low frequency inconsistent words. Pseudowords display a horizontal pattern, indicating that accuracy is primarily affected by phonological lesions.

A second linear mixed effects model examined how model phonological and semantic lesions influenced lexicality effects (Supplementary Table 6; Supplementary Fig. 7). We identified main effects of word type (β = 0.24, *t*(95) = 11.38, *p* < .001) and phonological lesion severity (β = -1.08, *t*(159) = -28.20, *p* < .001), such that pseudowords were read less accurately than real words and larger phonological lesions decreased reading accuracy of both real and pseudowords (Supplementary Fig. 6). Two two-way interactions were also found. There was an interaction of phonological lesion severity and word type (β = 0.07, *t*(294) = 11.95, *p* < .001), such that phonological lesions disproportionately decreased pseudoword reading accuracy (β = 0.07, *t*(294) = 11.95, *p* < .001). There was also an interaction of semantic lesion severity and word type (β = -0.19, *t*(95) = -4.62, *p* < .001), such that semantic lesions disproportionately reduce accuracy for words compared to pseudowords. In all, these models confirm that phonological model lesions impact both words and pseudowords strongly, but has a larger impact on pseudoword reading. Semantic model lesions instead disproportionately affect real word reading (Fig. 5).

### Effects of model lesions on semantic access during reading

While not the main focus of the current study, we also examined how the models’ semantic output in the reading task degrades with lesions to the phonological and semantic layers. The accuracy of semantic output is a metric of incidental semantic processing during oral reading. As expected, semantic output accuracy was primarily affected by semantic damage, such that models with small semantic lesions maintained a high level of semantic accuracy even in the presence of severe phonological damage (Fig. 6, upper left quadrant). However, in the presence of severe semantic damage, even perfectly intact phonology did not rescue semantic performance (Fig. 6, lower right quadrant).

**Fig. 6.**
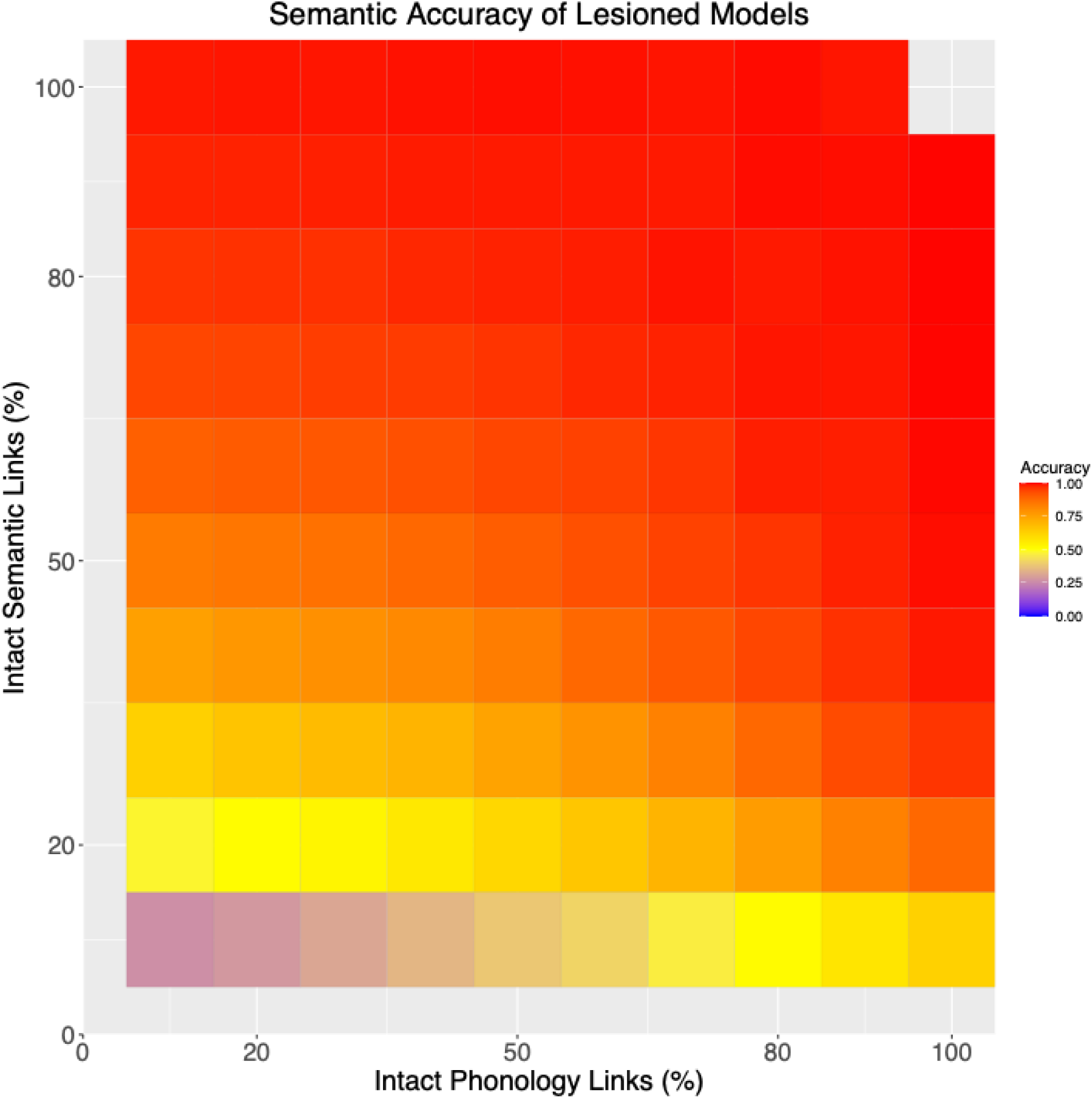
– Semantic output layer accuracy over lesioned models. The semantic accuracy of the model is shown as a function of lesion load in phonology and semantics. The asymmetric diagonal pattern shows that semantic output depends largely on the intactness of semantic links, but that with severe semantic lesions, intact phonological links can rescue semantic output to some degree. When semantics is intact, semantic output is accurate irrespective of phonological lesions (top row). In contrast, when semantics are lesioned, semantic accuracy depends on the intactness of phonology links (bottom four rows). With the most severe semantic lesions, even completely intact phonology cannot fully rescue semantic accuracy (bottom right).

### Assessing if individually matched lesioned models simulate real individual differences in alexic reading

Model reading performance and patient reading performance were highly correlated (all *r*’s 0.87 or higher, Fig. 7). Despite these high correlations, the matched models did differ in accuracy from stroke survivors. In paired t-tests, only pseudoword and low-frequency inconsistent word reading were not significantly different between the stroke survivors’ and matched-model (Table 5). High frequency words were overestimated by the largest amount (HFC: (*t*(82) = 8.1, *p* < .001; mean difference = 11.8; HFI: (*t*(82) = 8.5, *p* < .001; mean difference = 12.9)), showing that the lesioned models have a weaker frequency effect than is found in stroke survivors.

**Fig 7.**
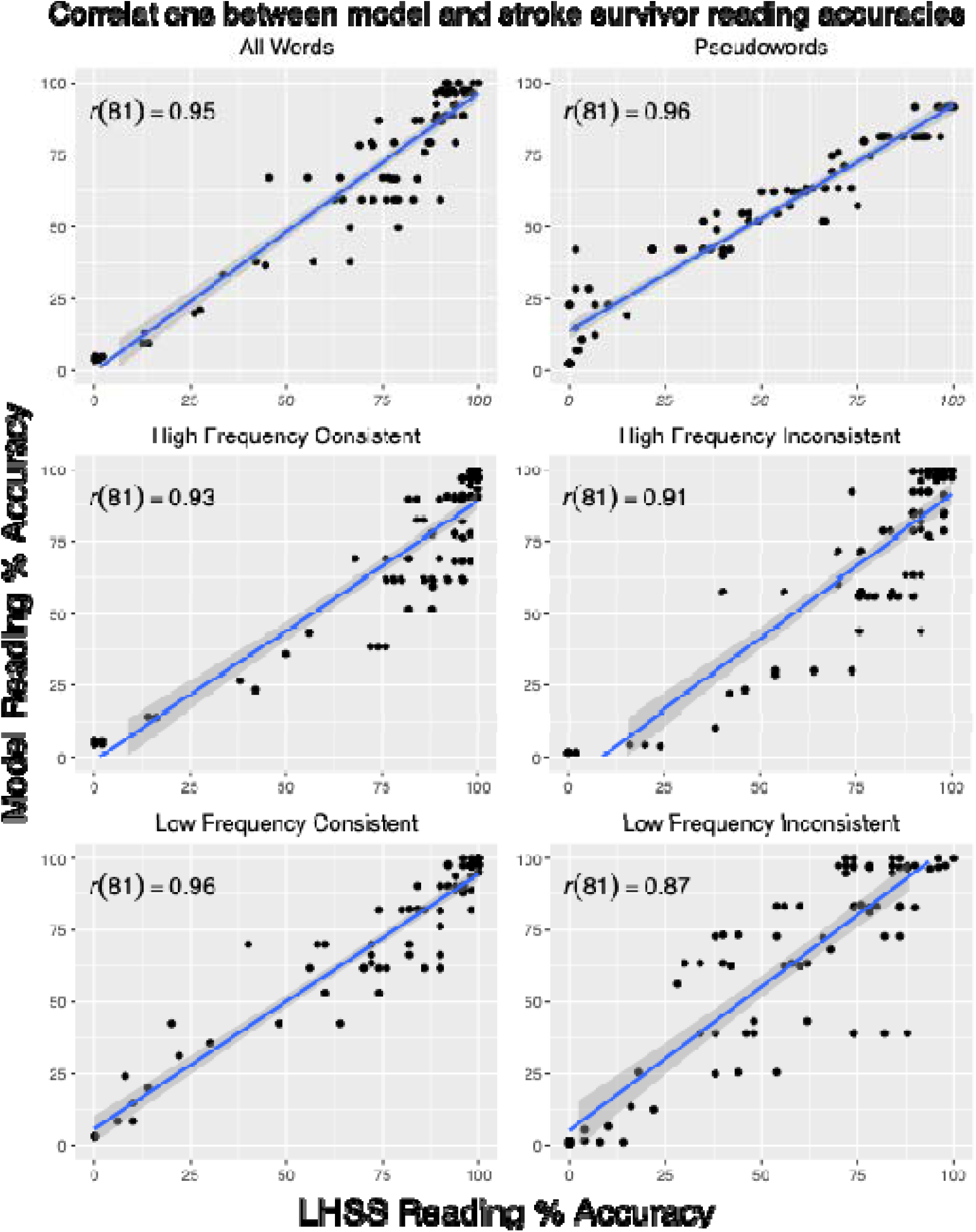
– Correlations between model and stroke survivor accuracy across word types. LHHS = left hemisphere stroke survivors. These plots assess if the matched models simulated the pattern of reading by word type that was observed in the stroke survivors.

**Table 5.**
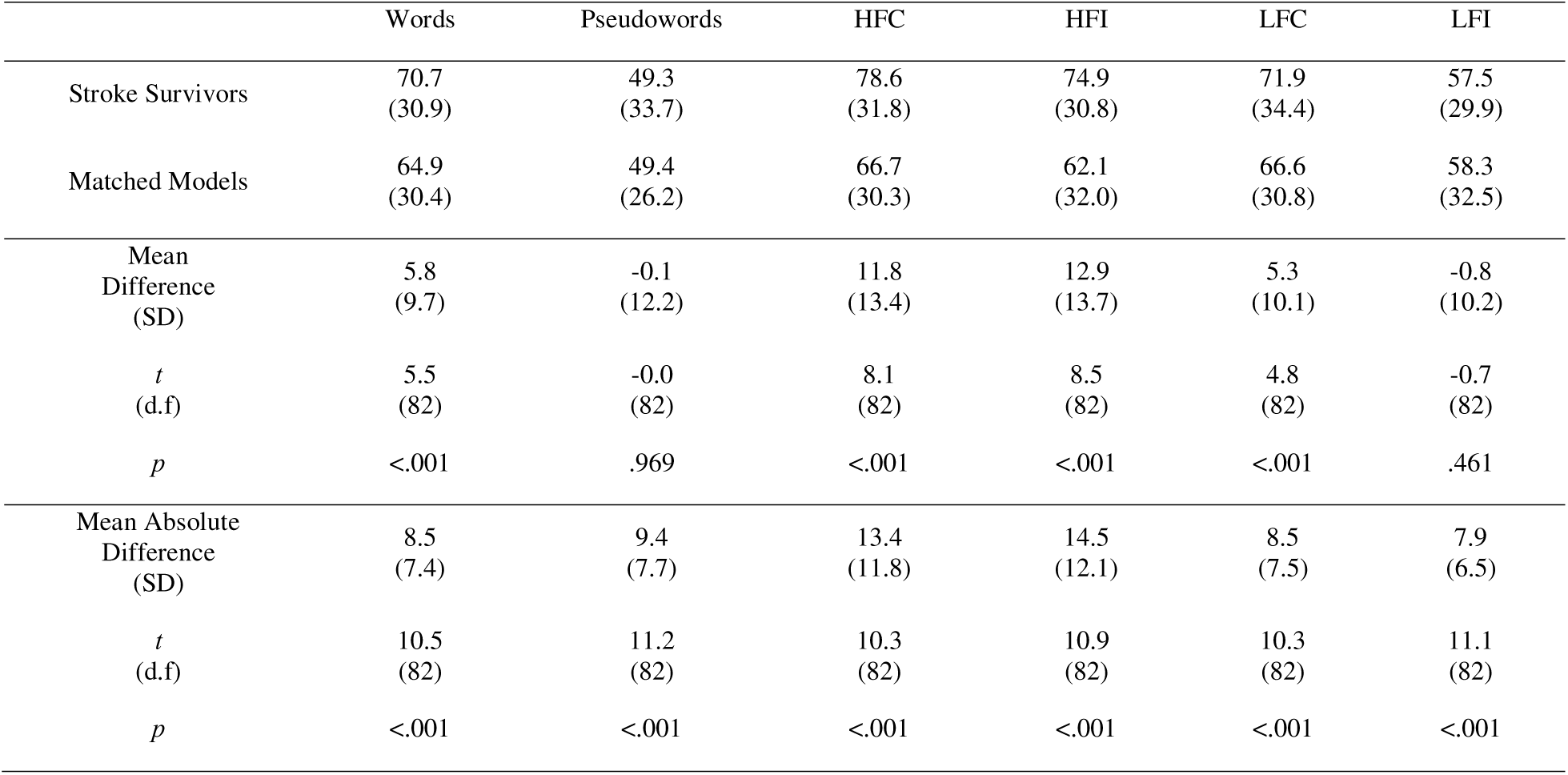
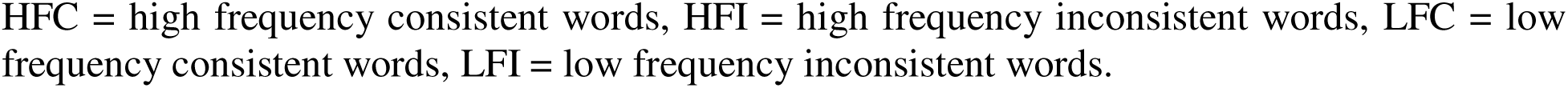
Stroke survivor and matched model reading.

|  | Words | Pseudowords | HFC | HFI | LFC | LFI |
| --- | --- | --- | --- | --- | --- | --- |
| Stroke Survivors | 70.7<br>(30.9) | 49.3<br>(33.7) | 78.6<br>(31.8) | 74.9<br>(30.8) | 71.9<br>(34.4) | 57.5<br>(29.9) |
| Matched Models | 64.9<br>(30.4) | 49.4<br>(26.2) | 66.7<br>(30.3) | 62.1<br>(32.0) | 66.6<br>(30.8) | 58.3<br>(32.5) |
| Mean Difference<br>(SD) | 5.8<br>(9.7) | -0.1<br>(12.2) | 11.8<br>(13.4) | 12.9<br>(13.7) | 5.3<br>(10.1) | -0.8<br>(10.2) |
| $t$<br>(d.f) | 5.5<br>(82) | -0.0<br>(82) | 8.1<br>(82) | 8.5<br>(82) | 4.8<br>(82) | -0.7<br>(82) |
| $p$ | <.001 | .969 | <.001 | <.001 | <.001 | .461 |
| Mean Absolute<br>Difference<br>(SD) | 8.5<br>(7.4) | 9.4<br>(7.7) | 13.4<br>(11.8) | 14.5<br>(12.1) | 8.5<br>(7.5) | 7.9<br>(6.5) |
| $t$<br>(d.f) | 10.5<br>(82) | 11.2<br>(82) | 10.3<br>(82) | 10.9<br>(82) | 10.3<br>(82) | 11.1<br>(82) |
| $p$ | <.001 | <.001 | <.001 | <.001 | <.001 | <.001 |

HFC = high frequency consistent words, HFI = high frequency inconsistent words, LFC = low frequency consistent words, LFI = low frequency inconsistent words.

Like the directly-measured composite scores, the matched model phonology and semantic scores were highly correlated (*r* = 0.89, *p* < .001, Fig. 4b). This shows that the matched models also capture the mixed impairment pattern found in the stroke survivors. The phonology composite scores and matched-model phonology scores were highly correlated (Spearman’s *r* = 0.77, *p* < .001, Fig. 8a) as were the semantic composite scores and matched-model semantic scores (Spearman’s *r* = 0.76, *p* < .001, Fig. 8b).

**Fig. 8.**
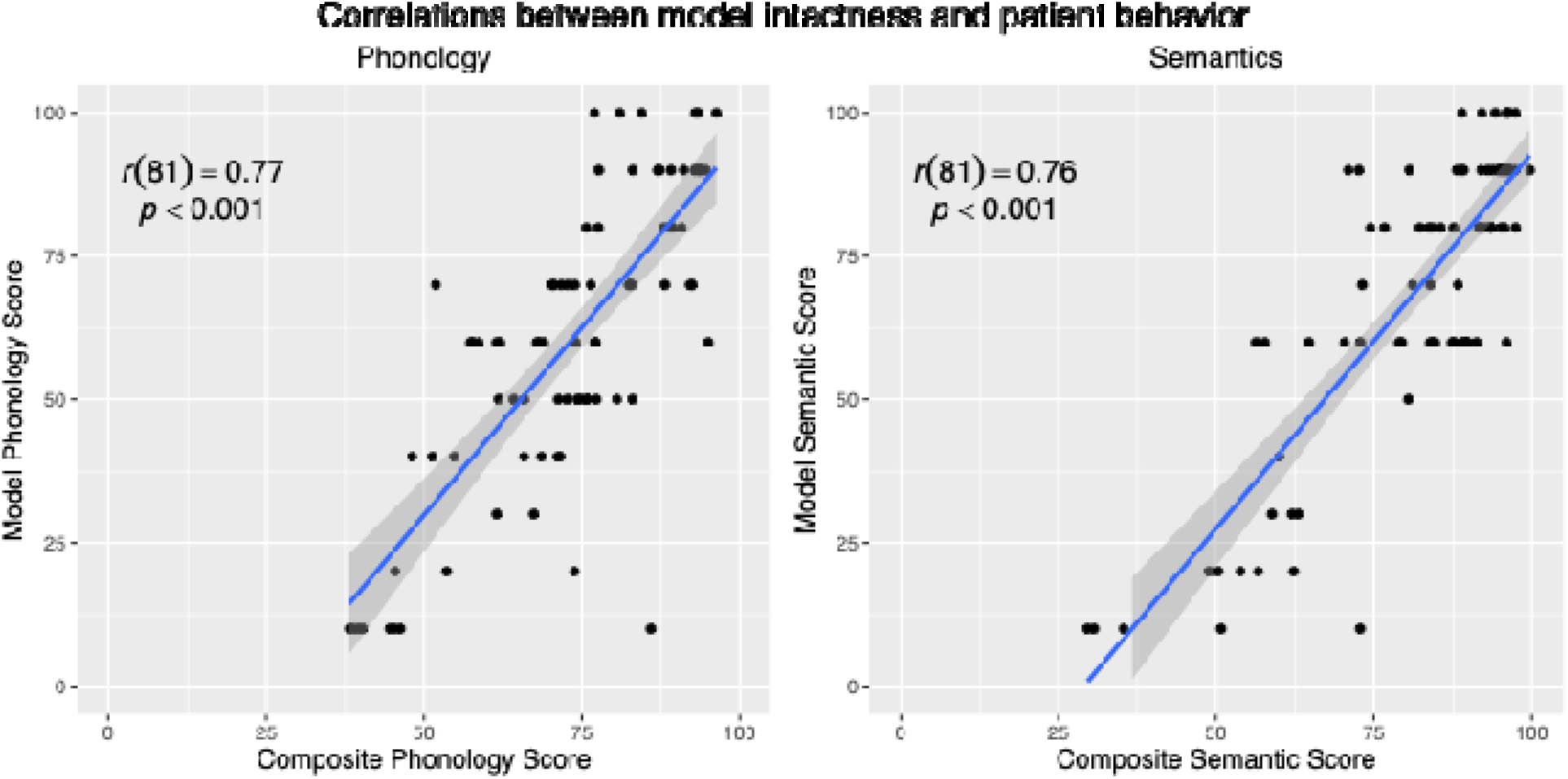
– a) Correlation between stroke survivor composite phonology scores and model phonology scores. b) Correlation between stroke survivor composite semantic scores and model semantic scores.

### Assessing factors that influence model fits

As shown in Fig. 7, the computational models were able to capture some word types, and some participants, better than others. We estimated a linear mixed effect model to understand which personal characteristics influenced how well the matched models predicted participants’ reading and how they interreacted with word types. Our critical tests are of the conditional marginal mean deviance for each word type against zero, which is the point at which the model would perfectly match a stroke survivors’ performance. Full parameter estimates for the model are in Supplementary Table 7. Analysis of marginal means showed that, at the mean phonological and semantic composite score values, all word types were significantly different from zero. Matched models underestimated reading accuracy on high frequency words by more than 10% (high frequency consistent words: Mean deviance=12.56, SE=1.64, *t*(167)=7.67, *p* < 0.001; high frequency inconsistent words: Mean deviance=12.89, SE=1.64, *t*(167)=7.87, *p* < 0.001). In contrast, deviations were less than 5% for low frequency words: the models modestly underestimated reading accuracy on low frequency consistent words (Mean deviance=3.25, SE=1.64, *t*(167)=1.99, *p* < 0.049) and modestly overestimated accuracy on low frequency inconsistent words (Mean deviance=-4.29, SE=1.64, *t*(167)=-2.62, *p* < 0.010).

The model identified three-way interactions of word type with semantic and phonology impairments, so we next assessed the deviance of the marginal means for each word type at +/- 1 SD for the phonological and semantic composite scores (Table 6, Figure 9b). Overall, the results suggested that 1) variation in stroke survivors’ phonological and semantic processing ability considerably affected model fit for high frequency words, but had relatively little impact on fit for low frequency words, 2) the models underestimated high frequency word reading most severely when phonology was impaired but semantics was intact.

**Fig. 9.**
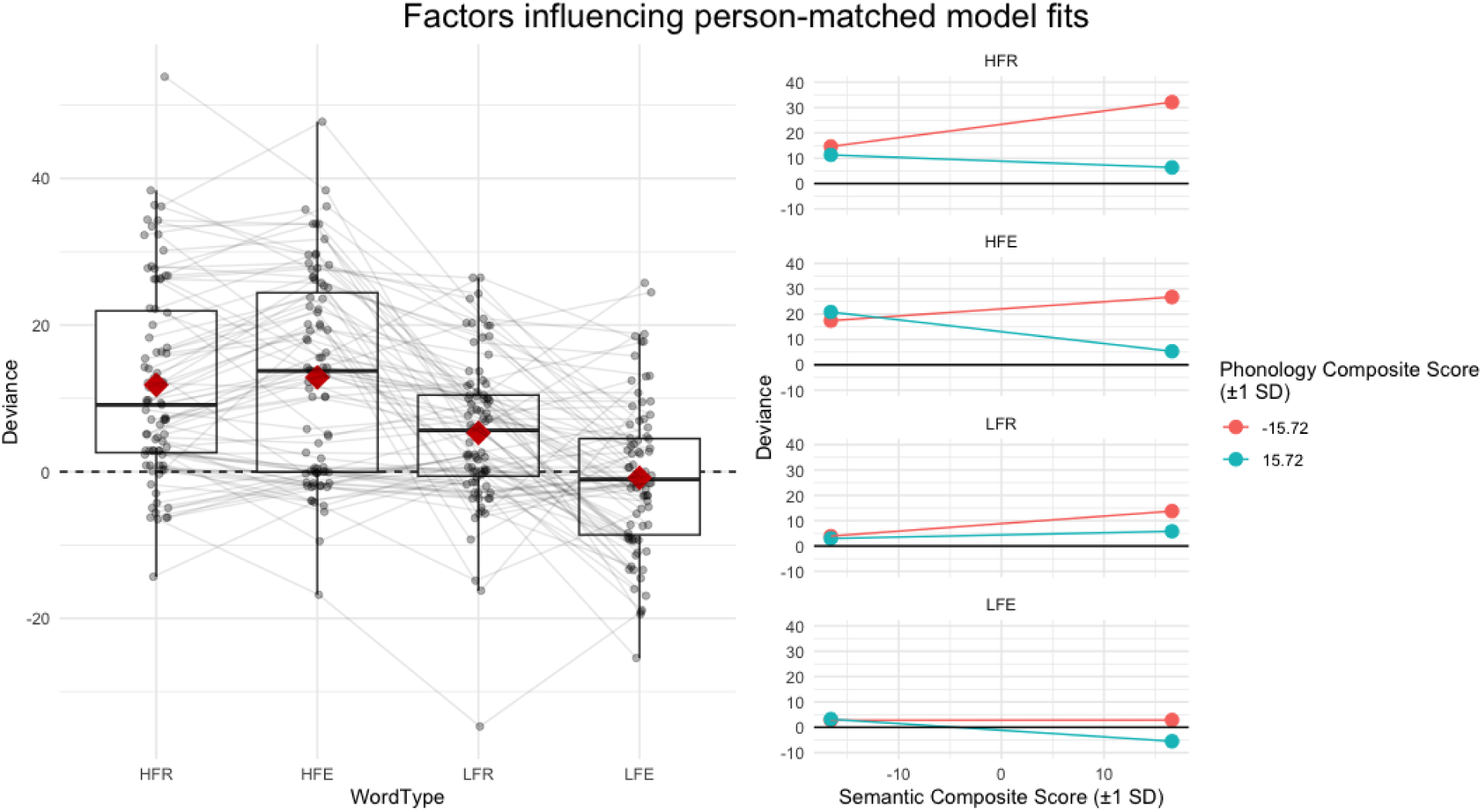
– Identifying the factors that influence how well the matched models fit their matched stroke survivor. a) Deviance (participant accuracy – matched-model accuracy in percent correct) by word type. Dots connected by lines indicate individual stroke survivors. Red diamonds indicate mean deviance. The dotted line at Y=0 indicates the point where the model prediction is exactly equal to stroke survivor reading accuracy. Positive values indicate cases where a stroke survivor was more accurate than their matched model. Horizontal jitter added to disambiguate overlapping points. b) Mixed-effects model estimates of the effects of semantic and phonological composite scores on deviance scores by word type.

**Table 6.** – Testing estimated marginal means against 0 for deviation between matched models and stroke survivors.

|  | Word Type | Marginal Mean | SE | df | *t* | *p* |
| --- | --- | --- | --- | --- | --- | --- |
|  | **HFC** | **14.63** | **1.81** | **161** | **8.10** | **< .001** |
| -1 SD Sem, | **HFI** | **17.43** | **1.81** | **161** | **9.65** | **< .001** |
| -1 SD Phon | **LFC** | **3.93** | **1.81** | **161** | **2.18** | **.042** |
|  | LFI | 2.79 | 1.81 | 161 | 1.55 | .124 |
|  | **HFC** | **32.14** | **3.60** | **159** | **8.92** | **< .001** |
| +1 SD Sem, | **HFI** | **26.76** | **3.60** | **159** | **7.43** | **< .001** |
| -1 SD Phon | **LFC** | **13.77** | **3.60** | **159** | **3.83** | **< .001** |
|  | LFI | 2.87 | 3.60 | 159 | 0.80 | .428 |
|  | **HFC** | **11.33** | **4.70** | **156** | **2.41** | **.009** |
| -1 SD Sem, | **HFI** | **20.84** | **4.70** | **156** | **4.43** | **<.001** |
| +1 SD Phon | LFC | 2.97 | 4.70 | 156 | 0.63 | .529 |
|  | LFI | 3.20 | 4.70 | 156 | 0.68 | .529 |
|  | **HFC** | **6.37** | **2.05** | **163** | **3.10** | **.009** |
| +1 SD Sem, | **HFI** | **5.34** | **2.05** | **163** | **2.60** | **.010** |
| +1 SD Phon | **LFC** | **5.81** | **2.05** | **163** | **2.83** | **.010** |
|  | **LFI** | **-5.51** | **2.05** | **163** | **-2.69** | **.010** |

HFC = high frequency consistent words, HFI = high frequency inconsistent words, LFC = low frequency consistent words, LFI = low frequency inconsistent words. Significant tests are bolded.

There were no significant effects of age, education, or apraxia severity. However, greater time since stroke was associated with greater underfitting (β = 0.07, *t*(75) = 3.18, *p* = .002).

## Discussion

Overall, the findings demonstrate that an ANN model with a classic triangle architecture can, to a large extent, simulate the individual variation in reading profiles of stroke survivors with alexia. Further, we found that the percentage of intact links in the matched models correlated with non-reading semantic and phonological composite scores. However, the stroke survivors did differ from their matched models in some systematic ways – primarily by underestimating the strength of the frequency effect. This and other differences between the model results and directly-measured results in the stroke survivors highlight where further optimization might improve the degree to which the models recapitulate real neurocognitive mechanisms of reading.

### Graded Process-Based Alexic Reading Deficits

Before discussing the main findings regarding the model fits, it is important to discuss the implications of the reading profiles in the stroke survivors. The interaction of group and consistency, such that stroke survivors were disproportionately poor at reading inconsistent words aloud compared to controls, is a hallmark of surface alexia (Coltheart et al., 1983) and often thought to be related to semantic impairments (Woollams et al., 2007). There was also a significant interaction of group and lexicality, where stroke survivors were disproportionately poor at reading pseudowords. This is the hallmark of phonological alexia (Beauvois & Derouesne, 1979; Derouesne & Beauvois, 1979), which reflects generalized phonological impairment (Welbourne & Lambon Ralph, 2007). The presence of both these effects across our sample suggests that traditional syndrome-based classification of alexia is insufficient to describe the reading impairments found in post-stroke alexia. Further, semantic and phonological processing impairments were highly correlated in the stroke survivors. If the stroke survivors showed a tendency towards the syndromic classifications, we would expect a small correlation or none at all. Given that continuous variation in reading performance was correlated with both semantic and phonological deficits, a graded process-based approach, focused on assessing generalized semantic, phonological, and visual processing (Crisp & Lambon Ralph, 2006), is likely to provide a better description of the underlying deficits in alexia. Using the graded process-based approach to better understand alexic deficits may lead to better diagnostic tools and better prediction of recovery and treatment effects.

Why are semantic and phonological deficits correlated in our sample, even when accounting for the overall size of stroke lesions? One possibility is that the anatomical proximity of dedicated semantic and phonological cortex makes it likely that any given perisylvian lesion impairs both. However, the correlation remained strong after controlling for lesion volume. An alternative explanation is that there is a general lexical processor underlying both semantic and phonological deficits that is often impacted by left hemisphere stroke. Functional, structural, and lesion evidence suggests that the pSTS may be a strong candidate for a convergence zone where orthographic, phonological, and semantic processes interact (Staples et al., 2025a; Wilson, Bautista, & McCarron, 2018). Because our non-reading language tasks broadly tapped lexical phonology and semantic processes, lesions to the pSTS may be responsible for impairing this lexical process.

### The Triangle Model Simulates Alexic Reading Performance Beyond Classic Syndromes

By allowing any given model to be damaged in both semantics and phonological capacity at the same time, we produced a range of possible reading deficits that the model could accommodate. We found that it is possible to simulate the oral reading of a large sample of stroke survivors by varying the severity of lesions to the phonological and semantic layers of model. Overall, the matched models fit their stroke survivor’s reading accuracies well. The correlations between the stroke survivors’ and matched models’ reading accuracies on the Real Word Test sets were uniformly high. The models thus capture the pattern of reading across stroke survivors. Further, although there were significant differences between the models’ numerical predictions of reading accuracy, the differences were generally small in magnitude across the group as a whole. These fits were obtained by matching on only two parameters: real word and pseudoword reading accuracy. Demonstrating that the models predict reading of the Real Word Test sets, which vary on frequency and consistency, from a fit obtained from just two parameters, indicates the strength of artificial neural networks as models of reading.

The pattern of results supports previous primary systems explanations of alexia: pseudoword reading was primarily affected by phonological lesions (Welbourne & Lambon Ralph, 2005, 2007) and inconsistent word reading was disproportionately affected by semantic lesions as compared to consistent word reading (Plaut et al., 1996; Woollams et al., 2007). Prior studies have demonstrated that computational models of reading can recapitulate idealized patterns associated with phonological and surface alexia, but the present results substantially extend this prior work to show that the triangle model can also simulate real-world variation in reading performance within and between idealized syndromic patterns.

The degree of success we found in modeling alexic reading behavior is remarkable considering that we only used graded lesions of two model components, the phonological and semantic layers. In reality, strokes can cause a wider variety of disruptions to the reading architecture, including disconnections of the central reading processors from input or output, lesions to orthographic processors, or disconnections between orthography, phonology, and semantics. We elected not to model all possible lesion configurations to avoid overfitting. The findings demonstrate that most of the variance in reading behavior in alexia can be accounted for only by damage to phonology and/or semantics. Future studies could use a hypothesis-driven lesion-matching approach to match more precise lesion configurations to individual people with alexia.

### Discrepancies Between Model Performance and Alexic Reading

Importantly, the matched models did deviate in reading ability from the stroke survivors. The most critical difference was that the matched models predicted worse high-frequency word reading than the stroke survivors displayed. This difference was modulated by the intactness of semantic and phonological processing in the stroke survivors. We consider two families of factors that may cause these deviations: parameters of the implemented models that may benefit from optimization, and the architecture of the model, where changes may improve correspondence between models and stroke survivors.

We have already discussed some implementation choices that might limit model fits, including only simulating lesions to two model components and matching models to participants based only on pseudoword and word reading. Other details of the model parameters, training, or the lesions that we implemented likely influenced how well they were capable of simulating the stroke survivors’ reading. For example, individuals who were tested longer after stroke were more underfit than those tested within the first few years of stroke. This result is to be expected because we chose not to vary the length of the Recovery period of training to keep the model instances maximally comparable. As such, the models do not account for ongoing recovery of reading ability over the course of years during the chronic period.

The most notable discrepancy in the model fits was the systematic underestimation of high frequency word reading. This may have resulted from training with a logarithmic frequency compression. While this approach is common in neural network models to reduce training time (Chang et al., 2024; Plaut et al., 1996), it may influence the magnitude of the frequency effect. Namely, it may cause the model to be exposed to low-frequency words more often than people are, permitting better learning. Alternatively, the fact that the models and stroke survivors were tested on different sets of words (100 words per frequency/consistency category for stroke survivors; 24 or 48 per category for the models) may have influenced this outcome. It cannot be ruled out, however, that neural network models do not demonstrate strong enough frequency effects across the range of possible lesions. The relatively weak semantic contributions identified by the deviance scores may contribute to the small frequency effect, as whole-word frequency is a large driver of semantic processing in models (Harm & Seidenberg, 2004). Our analysis of how semantic and phonological abilities affects model fit supports this explanation: for persons with poor phonological processing but good semantic processing, the models underestimated reading of high frequency words by a greater amount than for those with poor semantic and phonological processing. In contrast, high frequency words were much better fit when phonological processing was good.

These findings point towards the models having an imbalance in the division of labor between semantic and phonological processing: the phonological pathway is more sensitive to frequency than the semantic pathway. This may occur because the mappings from orthography to phonology are more systematic than between semantics and either orthography or phonology, making them easier for the model to learn. A systematic test of the strength of frequency in each pathway in the healthy model could be achieved by adopting a method from Harm & Seidenberg (2004), where accuracy is measured after a complete lesion of either the orthography-to-phonology connections or the semantics-to-phonology connections. Performing the same total-lesion method after a period of retraining following parametrically modulated lesions to semantics or phonology would assess how frequency effects change over the course of recovery. Further research should examine if these differences in model parameters affect the magnitude of the frequency effect.

Another, more interesting possibility to explain the discrepancies between model reading and alexic reading is that the model is missing critical components of the larger cognitive architecture present in humans. For example, a common sequalae of left-hemisphere stroke is AoS, which can be impaired independently of phonological or semantic processing. We excluded persons with severe apraxia from our sample to reduce the impact this would have on model fits. However, 29 stroke survivors (38%) in our sample had mild-to-moderate AoS. Directly modeling the motor planning and articulatory process would account for variance due to motor speech deficits, likely allowing for more accurate simulation of reading performance and better correspondence between model lesions and the underlying semantic and phonological processes. A second architectural consideration, motivated by neurobiological evidence (Staples et al., 2025b; Wilson et al., 2018), is the idea of a general lexical processor. Similar to ‘hub-and-spoke’ models of semantic processing (Dilkina, McClelland, & Plaut, 2010; Lambon-Ralph, Jefferies, Patterson, & Rogers, 2017; Rogers et al., 2004) in which modal semantic spokes are attached to an amodal integrative hub, this general lexical processor would have bidirectional connections to the orthographic, phonological, and semantic layers. This lexical processor would also serve to increase the sensitivity of the model to word frequency. Above, we show that the division of labor in the model differs from that in our participants. The relative ease of learning the systematic orthography-to-phonology mappings may interact with the architecture of the current model to produce this aberrant division of labor. Because the phonology-to-semantic connections are trained during the Preliterate phase, the easiest way to activate correct semantic representations during the Reading phase is via phonological mediation. The models quickly learn to activate the correct phonological output, which can then activate the correct semantic output via the pretrained connections, without having to learn strong orthography-semantic mappings. The addition of a lexical processor into the model allows a second route, lexical rather than phonological, by which semantics may be accessed.

### Limitations

In addition to the limitations discussed above, some general limitations about computational modelling of cognition are important to discuss. First, computational models generally have vast parameter spaces. Altering parameters without theoretical motivation could allow for any arbitrary behavior or pattern of behaviors to be simulated. Here, we have based model parameters on previous implementations of the triangle model and limited ourselves to actively manipulating only two parameters that we feel are warranted by previous research. However, other choices such as the learning rate or weight decay parameters, which have less apparent theoretical motivation, could alter the obtained fits.

Second, as suggested by the existence of competing models with different underlying mechanisms, many of which perform equivalently in predicting human reading behavior, producing a simulation of human behavior is not a guarantee that the model performs tasks like a human (Coltheart et al., 2001; Perry et al., 2007; Welbourne et al., 2011). Additional analyses determining if some aspects of the model generalize beyond the targeted behavior, such as assessing representational similarity between models and brains (Kriegeskorte, Mur, & Bandettini, 2008), could provide converging evidence that models are “human-like”, but more definitive assessment of human-like task performance is challenging to assess.

Beyond the limitations of modelling approaches, many stroke survivors receive speech-language therapy (SLT) during recovery. SLT may impart explicit reading strategies to stroke survivors, or provide explicit training in phonological (Brookshire, Conway, Pompon, Oelke, & Kendall, 2014) or semantic processing (Efstratiadou, Papathanasiou, Holland, Archonti, & Hilari, 2018). Neither differing strategies nor specific training for phonology or semantics are implemented in the model.

### Future Directions

Establishing the feasibility of large-scale modelling of alexia enables several areas of future research. The clear next step is to examine how changes to the lesions, the model’s architecture, and training might improve its capacity to fit the full range of post-stroke reading deficits. Beyond improving model fits, another next step is to examine if the activations of the model relate to the functional neural activations of stroke survivors performing language tasks in scanner, or if the magnitudes of the weights in different components of the models are related to functional and structural connectivity. Our results also have translational implications. Individualized computational models may provide a way to predict the efficacy of behavioral speech-language or neurostimulation interventions. For example, a variety of established treatments could be simulated on the matched model for a given stroke survivor. The best-performing treatment could then be deployed to optimize recovery.

## Conclusions

By matching lesioned computational models of reading to individual stroke survivors, we show that the models capture oral reading accuracy and recapitulate directly measured phonological and semantic processing ability. Deviances between the matched-models and stroke survivors suggest several ways in which the models could be improved, in particular the need to ensure that frequency effects in reading are better simulated. Our results support the primary systems hypothesis of reading and lay a foundation for predicting therapeutic responses from individualized models of recovery.

## Acknowledgement

We wish to thank the individuals who contributed to data collection, in alphabetical order: J. Vivian Dickens, Elizabeth Dvorak, Davetrina Seles Gadson, Catherine “Trini” Kelly, Elizabeth Lacey, Sachi Paul, Sarah Snider, and Candace van der Stelt. This work was funded by NIDCD/NIH (R01DC014960 and R01DC020446 to PET, R00DC018828 to ATD, and T32DC019481 to RS) and NCATS/NIH (T32DC019481 to RS).

## Footnotes

1 While further training produces perfect accuracy on the training corpus, preliminary simulations found that this amount of training produced qualitatively similar results to our sample of age-matched controls – specifically, that low frequency inconsistent words are read somewhat less accurately than other word types.

