## Supplementary Materials for "Simulating the spectrum, not the syndrome: Large scale individualized modeling of oral reading in stroke aphasia"

Supplementary Table 1 - Linear mixed effect logistic model predicting word reading accuracy

| *Predictors* | *Odds Ratios* | *std. Error* | *CI [95%]* | *z-statistic* | *p* |
| --- | --- | --- | --- | --- | --- |
| **(Intercept)** | **1045.77** | **524.15** | **391.57 – 2792.96** | **13.87** | **<0.001** |
| **PatientStatus [LHSS]** | **0.01** | **0.00** | **0.00 – 0.03** | **-8.65** | **<0.001** |
| **Frequency [low]** | **0.24** | **0.11** | **0.09 – 0.60** | **-3.03** | **0.002** |
| **Consistency [inconsistent]** | **0.13** | **0.06** | **0.05 – 0.32** | **-4.45** | **<0.001** |
| Age | 1.45 | 0.28 | 0.99 – 2.13 | 1.89 | 0.059 |
| Education | 1.19 | 0.23 | 0.81 – 1.75 | 0.89 | 0.375 |
| PatientStatus [LHSS] × Frequency [low] | 1.75 | 0.76 | 0.74 – 4.12 | 1.28 | 0.202 |
| **PatientStatus [LHSS] × Consistency [inconsistent]** | **4.95** | **2.05** | **2.20 – 11.13** | **3.87** | **<0.001** |
| Frequency [low] × Consistency [inconsistent] | 0.74 | 0.40 | 0.26 – 2.13 | -0.56 | 0.578 |
| (PatientStatus [LHSS] × Frequency [low]) × Consistency [inconsistent] | 0.64 | 0.30 | 0.26 – 1.62 | -0.94 | 0.348 |
| **Random Effects** | | | | | |
| σ^2^ | 3.29 | | | | |
| τ_00_ _Item_ | 0.98 | | | | |
| τ_00_ _SubjectCode_ | 4.97 | | | | |
| ICC | 0.64 | | | | |
| N _Item_ | 200 | | | | |
| N _SubjectCode_ | 145 | | | | |
| Observations | 29000 | | | | |
| Marginal R^2^ / Conditional R^2^ | 0.342 / 0.765 | | | | |

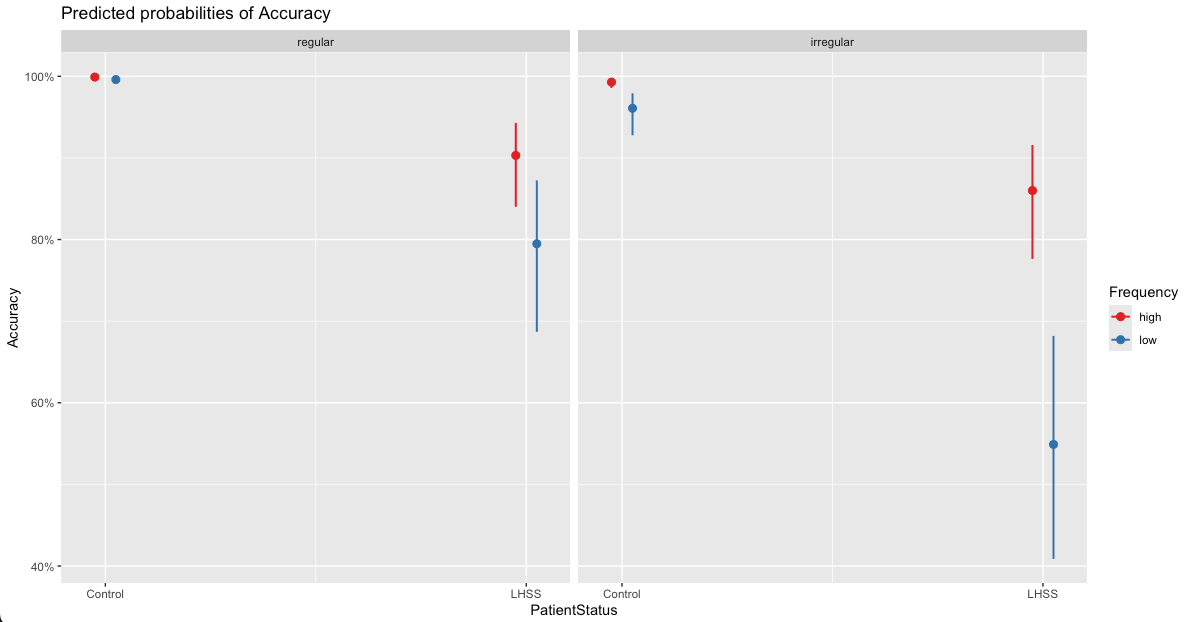

Supplementary Fig. 1 – Marginal means of a linear mixed effect model assessing the effect of patient status, frequency, and regularity on reading accuracy.

Supplementary Table 2 - Linear mixed effect logistic model predicting reading accuracy by lexicality

| *Predictors* | *Odds Ratios* | *std. Error* | *CI [95%]* | *z-statistic* | *p* |
| --- | --- | --- | --- | --- | --- |
| **(Intercept)** | **44.83** | **14.28** | **24.02 – 83.69** | **11.94** | **<0.001** |
| **PatientStatus [LHSS]** | **0.01** | **0.01** | **0.01 – 0.03** | **-11.09** | **<0.001** |
| **Lexicality [Real Word]** | **2.27** | **0.41** | **1.60 – 3.23** | **4.58** | **<0.001** |
| **PatientStatus [LHSS] × Lexicality [Real Word]** | **2.70** | **0.29** | **2.18 – 3.35** | **9.13** | **<0.001** |
| **Random Effects** | | | | | |
| σ^2^ | 3.29 | | | | |
| τ_00_ _Item_ | 1.32 | | | | |
| τ_00_ _SubjectCode_ | 4.97 | | | | |
| ICC | 0.66 | | | | |
| N _Item_ | 280 | | | | |
| N _SubjectCode_ | 145 | | | | |
| Observations | 38944 | | | | |
| Marginal R^2^ / Conditional R^2^ | 0.256 / 0.745 | | | | |

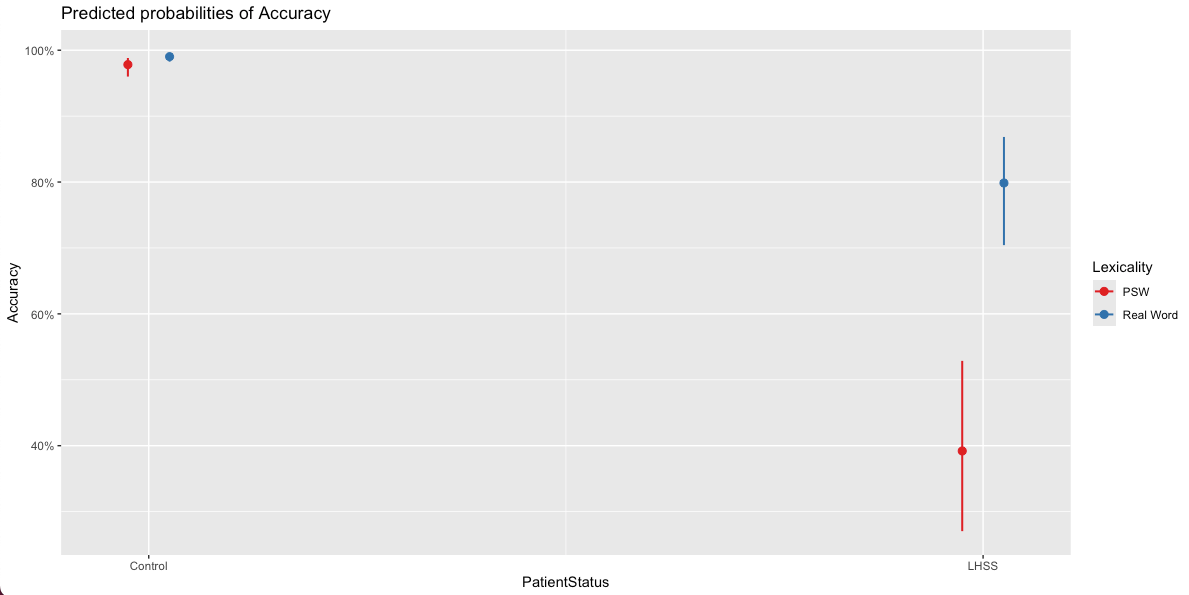

Supplementary Fig. 2 – Estimated marginal means from a linear mixed effect model assessing the effect of patient status and lexicality on reading accuracy.

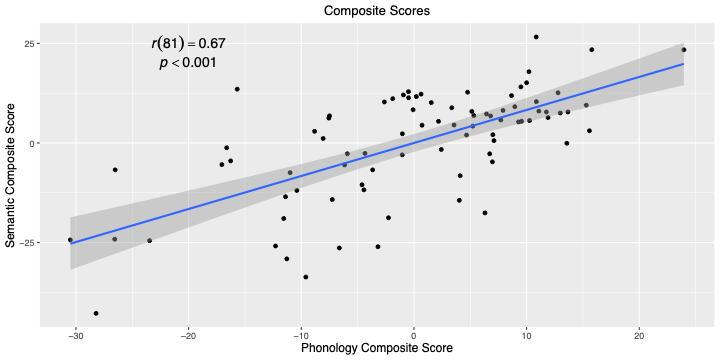

Supplementary Fig. 3 – Relationship between phonological and semantic composite scores after controlling for lesion volume, age, education, and ASRS AOS scores.

Supplementary Table 3 - Linear mixed effect logistic predicting ANN model word reading accuracy

| *Predictors* | *Estimates* | *std. Error* | *CI [95%]* | *t-statistic* | *p* | *df* |
| --- | --- | --- | --- | --- | --- | --- |
| **(Intercept)** | **0.52** | **0.03** | **0.45 – 0.59** | **15.04** | **<0.001** | **100.44** |
| **Frequency [low]** | **-0.03** | **0.01** | **-0.04 – -0.01** | **-4.37** | **<0.001** | **294.00** |
| **Consistency [consistent]** | **0.07** | **0.01** | **0.06 – 0.09** | **11.94** | **<0.001** | **294.00** |
| **Frequency [low] × Consistency [consistent]** | **0.02** | **0.01** | **0.00 – 0.04** | **2.22** | **0.027** | **294.00** |
| **Random Effects** | | | | | | |
| σ^2^ | 0.00 | | | | | |
| τ_00_ _Model_ | 0.12 | | | | | |
| ICC | 0.98 | | | | | |
| N _Model_ | 99 | | | | | |
| Observations | 396 | | | | | |
| Marginal R^2^ / Conditional R^2^ | 0.016 / 0.984 | | | | | |

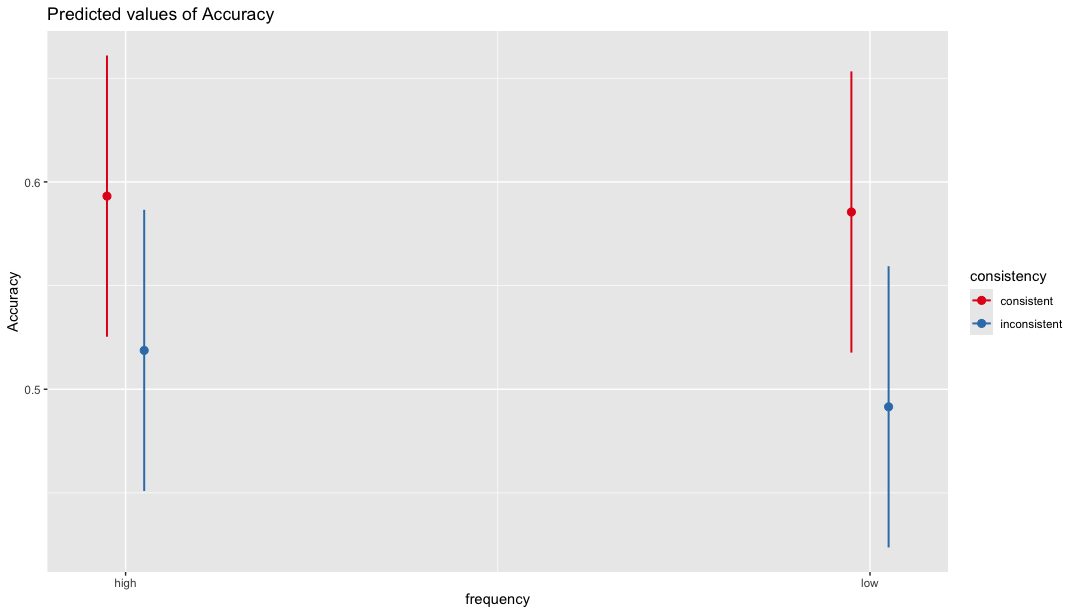

Supplementary Fig. 4 – Marginal means of a linear mixed effect model assessing the effect of frequency, and regularity across all lesioned model’s reading accuracy.

Supplementary Table 4 - Linear mixed effect model predicting ANN model reading accuracy by lexicality

| *Predictors* | *Estimates* | *std. Error* | *CI [95%]* | *t-statistic* | *p* | *df* |
| --- | --- | --- | --- | --- | --- | --- |
| (Intercept) | 0.57 | 0.03 | 0.51 – 0.64 | 17.50 | **<0.001** | 100.76 |
| WordType [PSW] | -0.12 | 0.01 | -0.13 – -0.10 | -15.41 | **<0.001** | 98.00 |
| **Random Effects** | | | | | | |
| σ^2^ | 0.00 | | | | | |
| τ_00_ _Model_ | 0.10 | | | | | |
| ICC | 0.97 | | | | | |
| N _Model_ | 99 | | | | | |
| Observations | 198 | | | | | |
| Marginal R^2^ / Conditional R^2^ | 0.032 / 0.973 | | | | | |

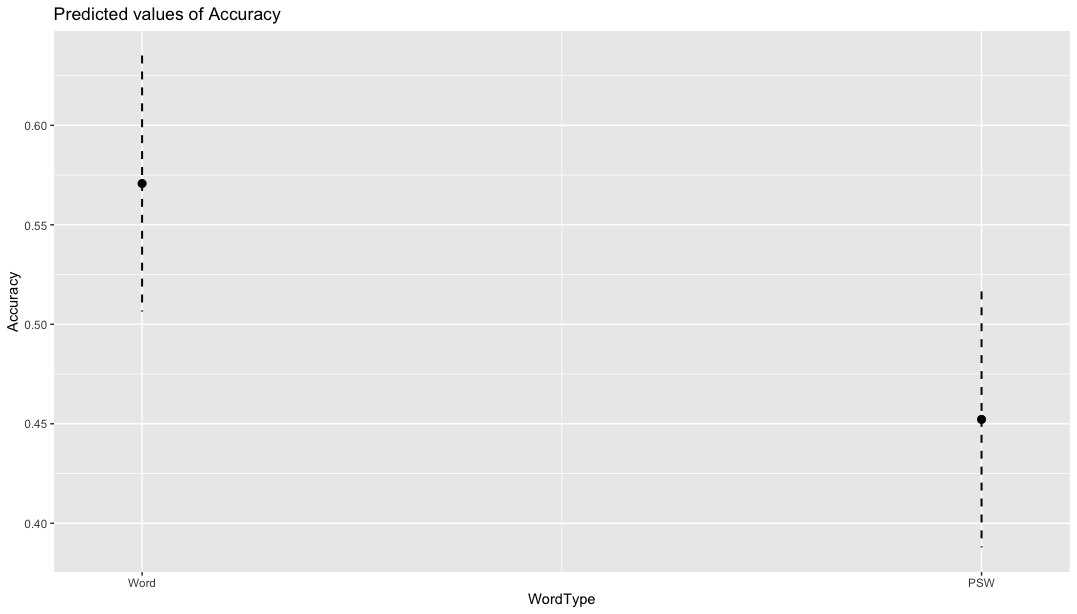

Supplementary Fig. 5 – Estimated marginal means of reading accuracy by lexicality across all lesioned models.

Supplementary Table 5 - Linear mixed effect model assessing how phonological and semantic lesions impact word reading in the ANN models

| *Predictors* | *Estimates* | *std. Error* | *CI [95%]* | *t-statistic* | *p* | *df* |
| --- | --- | --- | --- | --- | --- | --- |
| **(Intercept)** | **1.19** | **0.03** | **1.14 – 1.25** | **43.78** | **<0.001** | **125.46** |
| Frequency [low] | 0.01 | 0.02 | -0.02 – 0.04 | 0.57 | 0.567 | 285.00 |
| **Consistency [inconsistent]** | **0.04** | **0.02** | **0.01 – 0.07** | **2.32** | **0.021** | **285.00** |
| **Phon** | **-1.20** | **0.05** | **-1.30 – -1.10** | **-23.86** | **<0.001** | **125.46** |
| **Sem** | **-0.18** | **0.05** | **-0.28 – -0.08** | **-3.55** | **0.001** | **125.46** |
| Frequency [low] × Consistency [inconsistent] | -0.03 | 0.02 | -0.07 – 0.02 | -1.25 | 0.213 | 285.00 |
| Frequency [low] × Phon | -0.04 | 0.03 | -0.10 – 0.02 | -1.38 | 0.168 | 285.00 |
| **Consistency [inconsistent] × Phon** | **-0.06** | **0.03** | **-0.12 – -0.00** | **-2.13** | **0.034** | **285.00** |
| Frequency [low] × Sem | 0.00 | 0.03 | -0.06 – 0.06 | 0.00 | 0.998 | 285.00 |
| **Consistency [inconsistent] × Sem** | **-0.25** | **0.03** | **-0.30 – -0.19** | **-8.22** | **<0.001** | **285.00** |
| Phon × Sem | 0.12 | 0.09 | -0.06 – 0.30 | 1.28 | 0.202 | 125.46 |
| (Frequency [low] × Consistency [inconsistent]) × Phon | 0.01 | 0.04 | -0.07 – 0.10 | 0.34 | 0.735 | 285.00 |
| (Frequency [low] × Consistency [inconsistent]) × Sem | -0.02 | 0.04 | -0.10 – 0.06 | -0.49 | 0.626 | 285.00 |
| (Frequency [low] × Phon) × Sem | 0.01 | 0.06 | -0.10 – 0.12 | 0.16 | 0.873 | 285.00 |
| **(Consistency [inconsistent] × Phon) × Sem** | **0.14** | **0.06** | **0.03 – 0.25** | **2.53** | **0.012** | **285.00** |
| (Frequency [low] × Consistency [inconsistent] × Phon) × Sem | 0.06 | 0.08 | -0.10 – 0.21 | 0.75 | 0.453 | 285.00 |
| **Random Effects** | | | | | | |
| σ^2^ | 0.00 | | | | | |
| τ_00_ _Model_ | 0.00 | | | | | |
| ICC | 0.82 | | | | | |
| N _Model_ | 99 | | | | | |
| Observations | 396 | | | | | |
| Marginal R^2^ / Conditional R^2^ | 0.954 / 0.992 | | | | | |

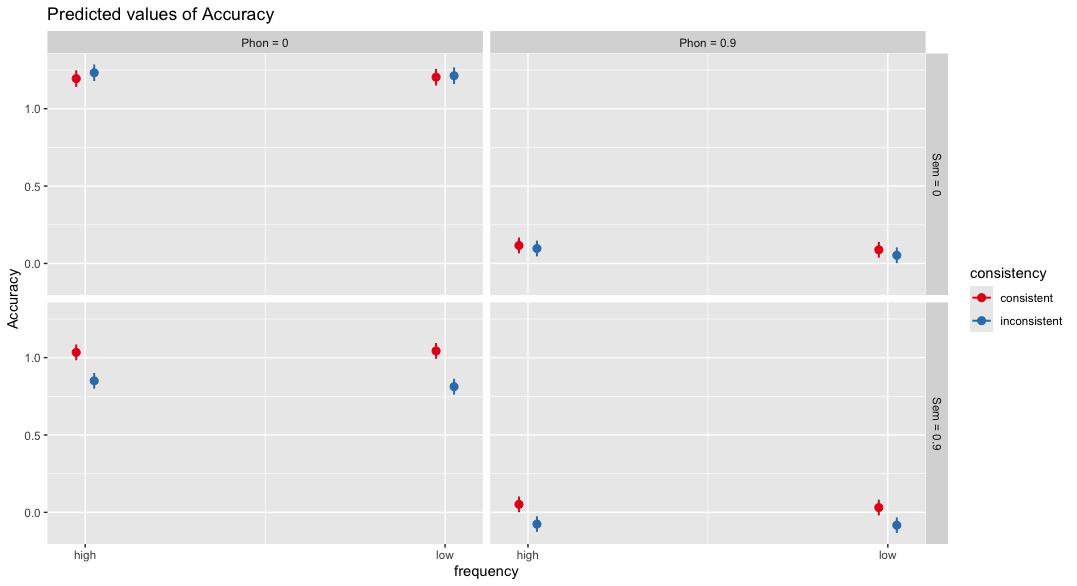

Supplementary Fig. 6 – Estimated marginal means of model semantic and phonological lesion severity on reading accuracy for high and low frequency and regularity words.

Supplementary Table 6 - Linear mixed effect model assessing how phonological and semantic lesions impact lexicality effects in the ANN models

| *Predictors* | *Estimates* | *std. Error* | *CI [95%]* | *t-statistic* | *p* | *df* |
| --- | --- | --- | --- | --- | --- | --- |
| **(Intercept)** | **0.94** | **0.02** | **0.90 – 0.98** | **45.43** | **<0.001** | **158.92** |
| **WordType [Words]** | **0.25** | **0.02** | **0.21 – 0.29** | **11.38** | **<0.001** | **95.00** |
| Sem | -0.01 | 0.04 | -0.09 – 0.06 | -0.37 | 0.712 | 158.92 |
| **Phon** | **-1.08** | **0.04** | **-1.15 – -1.00** | **-28.20** | **<0.001** | **158.92** |
| **WordType [Words] × Sem** | **-0.19** | **0.04** | **-0.27 – -0.11** | **-4.62** | **<0.001** | **95.00** |
| **WordType [Words] × Phon** | **-0.15** | **0.04** | **-0.23 – -0.07** | **-3.62** | **<0.001** | **95.00** |
| Sem × Phon | 0.03 | 0.07 | -0.11 – 0.17 | 0.41 | 0.685 | 158.92 |
| (WordType [Words] × Sem) × Phon | 0.10 | 0.07 | -0.05 – 0.25 | 1.34 | 0.184 | 95.00 |
| **Random Effects** | | | | | | |
| σ^2^ | 0.00 | | | | | |
| τ_00_ _Model_ | 0.00 | | | | | |
| ICC | 0.44 | | | | | |
| N _Model_ | 99 | | | | | |
| Observations | 198 | | | | | |
| Marginal R^2^ / Conditional R^2^ | 0.971 / 0.984 | | | | | |

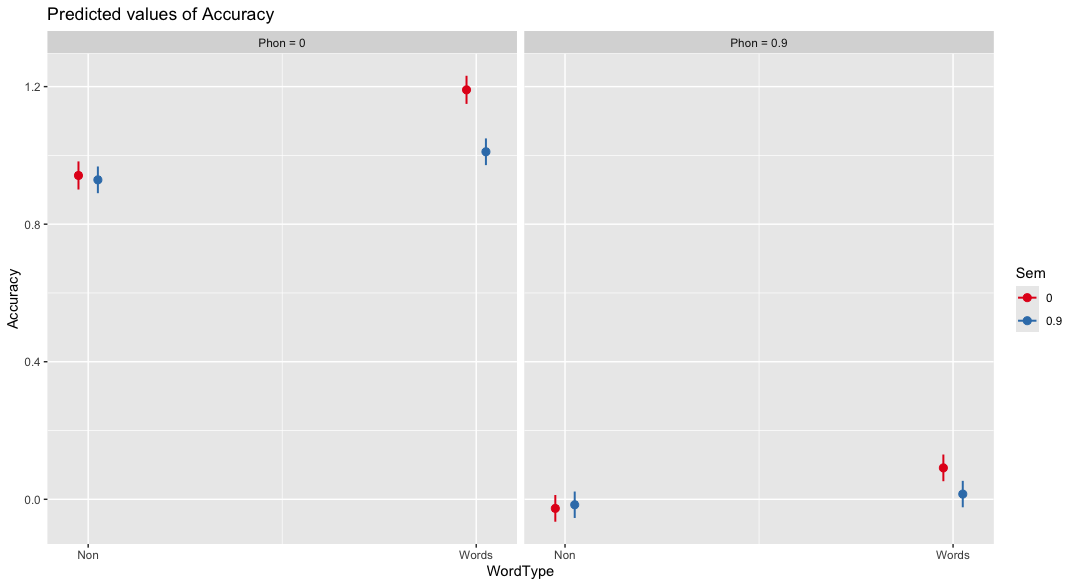

Supplementary Fig. 7 – Effects of model semantic and phonological reading severity on real and pseudoword reading accuracy.

Supplementary Table 7 – Estimates from a linear mixed-effects model predicting deviance between participant and matched-model reading accuracies by word type. These results are shown graphically in Fig. 8b.

|  | **Deviance (Participant Accuracy - Matched Model Accuracy)** | | | | | |
| --- | --- | --- | --- | --- | --- | --- |
| *Predictors* | *Estimates* | *std. Error* | *CI [95%]* | *t-statistic* | *p* | *df* |
| **(Intercept)** | **-100.70** | **26.22** | **-152.52 – -48.87** | **-3.84** | **<0.001** | **143.01** |
| WordType [HFE] | -0.46 | 23.26 | -46.29 – 45.36 | -0.02 | 0.984 | 237.00 |
| **WordType [LFR]** | **53.53** | **23.26** | **7.70 – 99.35** | **2.30** | **0.022** | **237.00** |
| **WordType [LFE]** | **72.10** | **23.26** | **26.27 – 117.92** | **3.10** | **0.002** | **237.00** |
| **AllSem** | **1.54** | **0.31** | **0.93 – 2.16** | **4.93** | **<0.001** | **164.41** |
| **AllPhon** | **1.21** | **0.44** | **0.35 – 2.08** | **2.78** | **0.006** | **158.58** |
| ASRS3 AOSSeverity | 1.46 | 1.29 | -1.10 – 4.02 | 1.13 | 0.260 | 75.00 |
| **Chronicity** | **0.07** | **0.02** | **0.03 – 0.11** | **3.18** | **0.002** | **75.00** |
| Age | 0.06 | 0.11 | -0.16 – 0.28 | 0.52 | 0.607 | 75.00 |
| Education | 0.28 | 0.39 | -0.50 – 1.07 | 0.73 | 0.469 | 75.00 |
| WordType [HFE] × AllSem | -0.04 | 0.30 | -0.64 – 0.56 | -0.13 | 0.896 | 237.00 |
| **WordType [LFR] × AllSem** | **-1.05** | **0.30** | **-1.65 – -0.45** | **-3.45** | **0.001** | **237.00** |
| **WordType [LFE] × AllSem** | **-1.33** | **0.30** | **-1.93 – -0.74** | **-4.39** | **<0.001** | **237.00** |
| WordType [HFE] × AllPhon | 0.30 | 0.42 | -0.52 – 1.12 | 0.73 | 0.467 | 237.00 |
| **WordType [LFR] × AllPhon** | **-0.84** | **0.42** | **-1.66 – -0.02** | **-2.02** | **0.045** | **237.00** |
| **WordType [LFE] × AllPhon** | **-0.88** | **0.42** | **-1.70 – -0.06** | **-2.12** | **0.035** | **237.00** |
| **AllSem × AllPhon** | **-0.02** | **0.00** | **-0.03 – -0.01** | **-3.87** | **<0.001** | **163.19** |
| (WordType [HFE] × AllSem) × AllPhon | -0.00 | 0.00 | -0.01 – 0.01 | -0.64 | 0.526 | 237.00 |
| **(WordType [LFR] × AllSem) × AllPhon** | **0.01** | **0.00** | **0.00 – 0.02** | **2.95** | **0.004** | **237.00** |
| **(WordType [LFE] × AllSem) × AllPhon** | **0.01** | **0.00** | **0.00 – 0.02** | **2.96** | **0.003** | **237.00** |
| **Random Effects** | | | | | | |
| σ^2^ | 69.79 | | | | | |
| τ_00_ _SubjectCode_ | 73.45 | | | | | |
| ICC | 0.51 | | | | | |
| N _SubjectCode_ | 83 | | | | | |
| Observations | 332 | | | | | |
| Marginal R^2^ / Conditional R^2^ | 0.356 / 0.686 | | | | | |
